# Inhibitory Fc Receptor sets a time limit on macrophage response to IgG

**DOI:** 10.64898/2026.09.21.753242

**Authors:** Annalise Bond, Erika T Snyder, Andrew Manion, Catherine Hardy, Kenny Kieu, Maxwell Z Wilson, Enoch Yeung, Meghan A Morrissey

**Affiliations:** Molecular Cellular and Developmental Biology Department, University of California; Santa Barbara, Santa Barbara, CA, USA; Interdisciplinary Quantitative Biosciences Program, University of California; Santa Barbara, Santa Barbara, CA, USA; Department of Mechanical Engineering, University of California, Santa Barbara; Santa Barbara, CA, USA

**Keywords:** Fc Receptors, FcγRIIB, Phagocytosis, IgG, Inhibitory Receptors, Inflammation, Macrophage Signaling, Signal Integration

## Abstract

Antibodies engage both activating Fc Receptors and the inhibitory receptor FcγRIIB. Why macrophages need a dedicated inhibitory receptor rather than simply tuning activating receptor signaling is unclear. Using DNA-based chimeric receptors and in silico modeling, we independently controlled activating and inhibitory Fc Receptors. We found that FcγRIIB imposed a time limit on macrophage phagocytosis and ERK signaling. The time limit is due to activating Fc Receptors converting PI(4,5)P2 to PI(3,4,5)P3, which is subsequently converted to PI(3,4)P2 by FcγRIIB. This leads to a pulse of active signaling, which is sufficient for phagocytosis of small bacteria-sized targets but not phagocytosis of large targets and TNFα secretion. Unlike engaging FcγRIIB, reducing activating Fc Receptor signaling decreased initiation of phagocytosis, the speed of PI(3,4,5)P3 generation, and the amplitude of ERK signaling. Our results demonstrate that FcγRIIB controls the duration of IgG signaling, while the activating Fc Receptors control sensitivity.

## Introduction

IgG antibodies bind antigens, marking targets for destruction by immune cells (*1*). Therapeutic monoclonal antibodies are a common and versatile tool for controlling immune responses, acting as receptor blockades, signaling agonists or flagging pathogenic cells for immune destruction. Macrophages phagocytose IgG-bound targets to destroy virally infected cells, bacteria, or other pathogens. Antibody-dependent phagocytosis is a key mechanism of several therapeutic antibodies including rituximab (anti-CD20) and trastuzumab (anti-Her2) (*2*).

Immune cells recognize antibodies through Fc Receptors. Even antibodies originally designed as blockade antibodies require Fc Receptors for full efficacy (*3*, *4*). The Fc Receptor family includes multiple receptors that bind IgG with varied affinities (*5*). Most Fc Receptors signal through an intracellular immune tyrosine activating motif (ITAM) that recruits SYK kinase to activate downstream signaling. However, both humans and mice have one inhibitory Fc Receptor, FcγRIIB, with an intracellular immune tyrosine inhibitory motif (ITIM) (*6*). In B cells, FcγRIIB is the only Fc Receptor expressed and prevents B cells from reacting to an antibody-antigen complex that simultaneously binds the B cell receptor and an existing IgG (*7*). However, many immune cells, including macrophages, express both activating and inhibitory Fc Receptors (*5*). Myeloid-specific knockout of FcγRIIB leads to increased inflammation and susceptibility to inflammatory disorders like nephrotoxic nephritis and rheumatoid arthritis (*8–11*). Why macrophages require both an activating and inhibitory receptor recognizing the same ligand remains unclear.

FcγRIIB is proposed to limit macrophage sensitivity (*11*). However, macrophages could accomplish this by simply expressing less activator (*12*). Fc Receptor expression is quite plastic, and can be modified by exposure to different cytokines or across different macrophage subsets (*13*). In addition, the immune system produces multiple IgG isotypes that vary in affinity for the activating and inhibitory Fc Receptors (*14*). The amount of activating and inhibitory receptor engagement, known as the A:I ratio, is carefully considered when designing new antibody therapies (*14*, *15*). Despite this, there have been relatively few studies on how immune cells integrate activating and inhibitory Fc Receptor signaling.

Here, we asked how a dedicated inhibitory Fc Receptor differs from controlling macrophage responses through activating Fc Receptors alone. We found that activating Fc Receptors control macrophage sensitivity, determining whether signaling initiates, while FcγRIIB sets a time limit on how long signaling persists. Macrophages have several responses to IgG that occur on different time scales - phagocytosis is rapid while cytokine secretion requires sustained signaling. Our findings suggest a mechanism for immune cells to separately control sensitivity and output.

## Results

### Molecular control of FcγRIIB

Since IgG activates both FcγRIIB and activating Fc Receptors, prior studies could not selectively activate FcγRIIB. We previously used a DNA chimeric antigen receptor (CAR) to study activating Fc Receptors (*16*), and DNA CARs have uncovered many principles of T cell signaling (*17*, *18*). To isolate the effect of FcγRIIB, we designed an FcγRIIB DNA CAR activated by a complementary ssDNA instead of IgG **(Fig 1A, Table S1)**. The FcγRIIB DNA CAR contains the intracellular and transmembrane domains of the mouse FcγRIIB. The FcγRIIB extracellular domain is replaced by a Halo tag that covalently binds single stranded DNA (ssDNA) with a 5’-Halo Ligand. To determine if this FcγRIIB DNA CAR suppresses phagocytosis, we expressed the construct in the RAW264.7 macrophage-like cell line. As a phagocytic target, we used silica beads coated in supported lipid bilayers to mimic the fluid cell membrane. We functionalized beads with either anti-biotin IgG alone or IgG and a complementary ligand DNA, then measured phagocytosis. Adding ligand DNA suppressed phagocytosis by 50% **(Fig 1B, S1A,B)**, showing the FcγRIIB DNA CAR inhibits phagocytosis.

**Figure 1.**
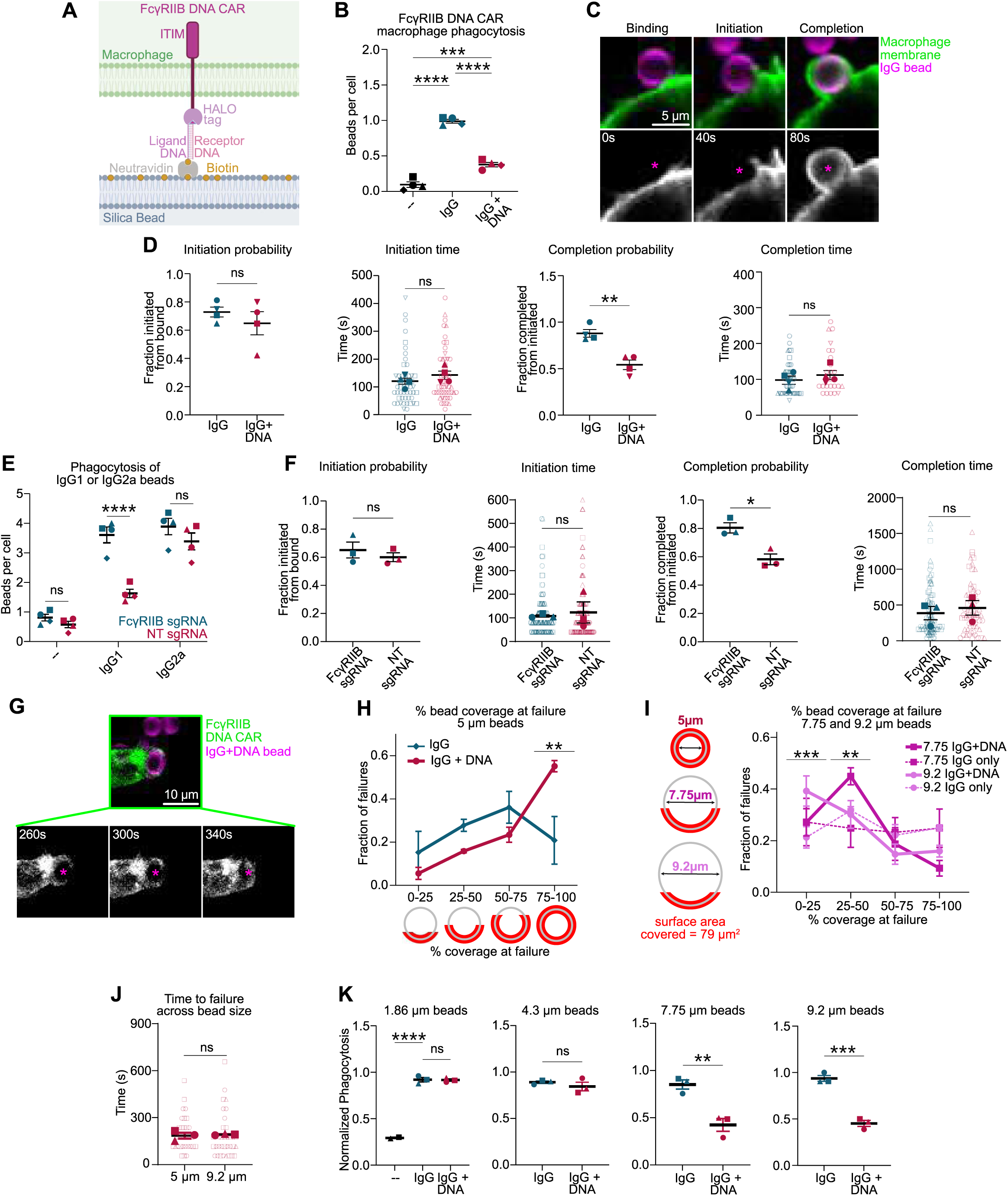
FcγRIIB sets a time limit for phagocytosis. **(A)** Schematic of DNA FcγRIIB CAR. **(B)** FcγRIIB DNA CAR RAW264.7 macrophages were incubated with supported lipid bilayer coated beads bound to neutravidin (--), IgG and neutravidin (IgG), or IgG, neutravidin and biotinylated ligand DNA (IgG+DNA). The average number of 5 µm diameter beads phagocytosed per macrophage was quantified by spinning disk confocal microscopy. **(C)** Timelapse confocal microscopy shows bead binding, initiation, and completion of phagocytosis by a RAW264.7 macrophage (mCh-CAAX, green top; greyscale below) encountering a supported lipid bilayer (atto647, magenta) coated bead bound to IgG. **(D)** Time lapse microscopy was used to quantify the fraction of beads bound to a macrophage that proceeded to initiation of phagocytosis; the time from binding to initiation; the fraction of beads that proceeded from initiation to completion; the time between initiation and completion of phagocytosis. **(E)** 9 µm diameter beads with 6xHis CD20 incorporated into their supported lipid bilayer were opsonized with anti-CD20 IgG1, IgG2a or no IgG (--), then incubated with HoxB8 macrophages infected with sgRNA targeting FcγRIIB or non-targeting (NT) sgRNA. **(F)** Timelapse microscopy was used to quantify the stages of phagocytosis as described in D for macrophages infected with a FcγRIIB or nontargeting sgRNA encountering 9 µm diameter IgG1 beads. **(G)** Images show a FcγRIIB DNA CAR (green; greyscale below) RAW264.7 macrophage failing to phagocytose a IgG+DNA bead (atto647, magenta; 5 µm diameter). **(H)** The percentage of the target covered by FcγRIIB DNA CAR macrophage membrane at failure for IgG or IgG+DNA targets was measured by timelapse microscopy. **(I)** Schematic highlights the same surface area (79 μm^2^) on beads with varying diameters. The graph shows the percentage of the target covered by the macrophage membrane at failure for IgG (solid) or IgG+DNA (dashed) targets for larger beads. **(J)** The time from initiation to failure of phagocytosis for 5 μm and 9.2 μm diameter beads was measured using timelapse microscopy. **(K)** FcγRIIB DNA CAR macrophages were incubated with beads of varying diameter containing IgG or IgG + DNA. In H and I, points represent the mean of 3 independent experiments. In other graphs, solid data points represent the mean of an independent experiment, while unfilled data points represent individual cell measurements. Data points from replicates performed on the same day are denoted by symbol shape. In all graphs, bars represent the mean +/- SEM. *indicates p<0.05, **indicates p<0.01, ***indicates p<0.001, ****indicates p<0.0001 by ordinary one-way ANOVA with Tukey’s multiple comparisons test (B); unpaired two-tailed t test (D, F, J, K); two-way ANOVA with Šídák’s multiple comparisons test (E, H, I); and ordinary one-way ANOVA with Dunnett’s multiple comparisons test (K, first graph).

In B cells, FcγRIIB functions via the 5’ inositol phosphatases SHIP1 and SHIP2 (*19–21*). PI(3,4,5)P3 (PIP3), a substrate of SHIP1, is a signaling lipid required for phagocytosis so dephosphorylating PIP3 could prevent phagocytosis (*22*). To test whether the FcγRIIB DNA CAR suppresses phagocytosis via inositol phosphatases, we inhibited SHIP1 and SHIP2, or SHP1 and SHP2 then measured phagocytosis of targets containing IgG or IgG and ligand DNA. Inhibiting inositol phosphatases SHIP1 and SHIP2 eliminated the inhibitory effect of the FcγRIIB DNA CAR **(Fig S1C)**. In contrast, inhibiting the tyrosine phosphatases SHP1 and SHP2 had no measurable effect, indicating the FcγRIIB DNA CAR inhibits through SHIP phosphatases.

### FcγRIIB inhibits completion of phagocytosis

Phagocytosis requires target binding, sufficient activating signals to initiate phagocytosis, coordinated extension of actin and membrane around the phagocytic target to form the cup, and membrane fusion to complete the phagosome (*16*, *23*, *24*) **(Fig 1C)**. We next asked how FcγRIIB affects these processes **(Fig 1D)**. FcγRIIB DNA CAR signaling did not significantly affect phagocytosis initiation frequency or the time between binding and initiation. However, for half of ssDNA-ligand targets, macrophages failed to complete phagocytosis, with the phagocytic cup retracting or stalling indefinitely. Previously, we showed the inhibitory ligand CD47 affects phagocytosis completion by preventing reaching phagocytosis (*25*). However, unlike CD47, FcγRIIB did not measurably impact phagocytosis speed or cup morphology **(Fig 1D, S1D).** Together, these data suggest FcγRIIB causes phagocytic failure without affecting preceding stages.

FcγRIIB has been proposed to raise the threshold for initiating a macrophage response, dampening macrophage sensitivity (*11*). However, our data suggested FcγRIIB regulates phagocytosis completion, not initiation. To directly test whether FcγRIIB decreases macrophage IgG sensitivity, we transduced HoxB8-driven, conditionally immortalized ROSA-Cas9 macrophages with an sgRNA targeting FcγRIIB **(Fig S2A,B)**. We then measured phagocytosis relative to control macrophages infected with a non-targeting sgRNA. IgG isotypes vary in affinity for the activating and inhibitory Fc Receptors. IgG1 has the highest affinity for FcγRIIB, while IgG2a has very low affinity for FcγRIIB (*14*, *26*). Eliminating FcγRIIB did not affect phagocytosis of IgG2a-opsonized beads and increased phagocytosis of IgG1 beads, as expected **(Fig 1E, S2C)**.

We next titrated IgG1 on the targets to determine if FcγRIIB lowered the density of IgG required to initiate phagocytosis. Like control macrophages, FcγRIIB sgRNA macrophages did not phagocytose at low IgG densities **(Fig S2D)**. At higher densities, both frequently phagocytosed, but FcγRIIB sgRNA macrophages phagocytosed more beads.

We used timelapse microscopy to determine if FcγRIIB sgRNA macrophages were more likely to initiate or complete phagocytosis. FcγRIIB sgRNA only increased the frequency of completion, as predicted by our FcγRIIB DNA CAR **(Fig 1F, S2E-G)**.

### FcγRIIB sets a time limit for phagocytosis

In our timelapse imaging of phagocytosis, we noted that phagocytosis often failed just before completion, when the macrophage membrane covered the majority of the target surface **(Fig 1G; Fig S2E)**. When the FcγRIIB DNA CAR was activated, most failures occurred when more than 75% of a 5 µm target was covered **(Fig 1H)**. In contrast, the rare phagocytic failures of IgG targets were evenly distributed across the target surface.

This data suggested two models for FcγRIIB function. First, FcγRIIB could inhibit a late step in phagocytosis such as membrane re-sealing during cup closure. Prior studies have implicated dynamin as an essential regulator of cup closure (*27*, *28*). We titrated a dynamin inhibitor (MiTMAB) to select a dose that suppressed phagocytosis by 50% like the FcγRIIB DNA CAR **(Fig. S3A)**. Macrophages treated with this inhibitor failed phagocytosis when the phagocytic cup had covered at least 75% of the target, suggesting that inhibiting cup closure could produce a similar phenotype to FcγRIIB **(Fig S3B-C)**.

The second model is that FcγRIIB sets a time limit for phagocytosis. In other biological circuits, one signal turns on both a positive and negative regulator of the same output (*29–31*). If the inhibitor is delayed relative to the activator, this creates an incoherent feed forward loop and a pulse of downstream signaling. Paired activating Fc Receptors and FcγRIIB could similarly generate a pulse of signaling.

To test if FcγRIIB inhibits phagocytosis at a specific stage or after a consistent time, we varied phagocytic target size. If FcγRIIB regulates a specific stage of phagocytosis, its activation will cause failure at the final stages regardless of target size. If it instead sets a time limit, phagocytosis will fail at a consistent time. Since larger targets take longer to engulf, phagocytosis would fail with a smaller fraction of the bead covered (*32*). We found as bead size increased, phagocytosis failed with protrusions covering a smaller fraction of the target area **(Fig 1I, S3D, Table S2)**. This fraction corresponded to a similar total surface area across all bead sizes. This is inconsistent with FcγRIIB regulating a specific step required to complete phagocytosis, like cup closure.

We next measured how long macrophages engaged with a target before that phagocytic cup stalled or retracted. FcγRIIB DNA CAR macrophages spent similar time engaging 9.2 µm and 5 µm diameter targets before stalling or retracting the cup **(Fig 1J)**. FcγRIIB DNA CAR activation failed to inhibit phagocytosis of beads that were small enough to be phagocytosed within the observed time limit **(Fig 1K)**. In addition, FcγRIIB sgRNA macrophages spent longer engaging with targets than control macrophages before phagocytosis failed **(Fig S2G)**. This is consistent with FcγRIIB regulating the duration of attempted phagocytosis.

### In silico model predicts that paired activating and inhibitory Fc Receptors generate a pulse of PIP3

We next sought to uncover the mechanism generating the FcγRIIB-dependent time limit. Since activating and inhibitory Fc Receptor pathways converge on PIP3, we modeled PIP3 kinetics after IgG binding in silico. The phagocytic cup is separated from the rest of the plasma membrane by a diffusion barrier that prevents lipids from entering or exiting (*33*, *34*). During phagocytosis, SYK activates phosphoinositol 3-kinase (PI3K), which converts PI(4,5)P2 into PIP3 (*35*). PIP3 recruits many downstream effectors and is essential for phagocytosis progression and completion (*22*). SHIP1/2 are 5’ inositol phosphatases that regulate phagocytosis by targeting PIP3, converting it to PI(3,4)P2 (*36*). PI(3,4)P2 is not converted back to PIP3 within the time of phagocytosis, since no 5’ inositol kinases act at the cup (*37*, *38*). We modeled these PIP dynamics using previously published enzyme kinetics **(Fig 2A, Table S3,4)**. When FcγRIIB is not active, PIP3 is generated until the precursor PI(4,5)P2 is depleted (*39*) **(Fig 2B)**. When FcγRIIB is active, PIP3 is initially generated more quickly than it is depleted. As PIP3 accumulates and the precursor PI(4,5)P2 is depleted, PIP3 degradation begins to outpace synthesis. This produces a local maximum of PIP3 concentration followed by a rapid decay **(Fig 2C**). We also built a minimalistic, unitless model where the activating and inhibitory arms are represented by a single variable. This simplified system also generated a transient pulse of PIP3 **(Fig. S4A-B; Table S5)**. Together, these models suggest that activating Fc Receptors generate PIP3, which the inhibitory FcγRIIB subsequently depletes, leading to a pulse of signaling that terminates when PI(4,5)P2 is flushed from the signaling compartment (*40*).

**Figure 2.**
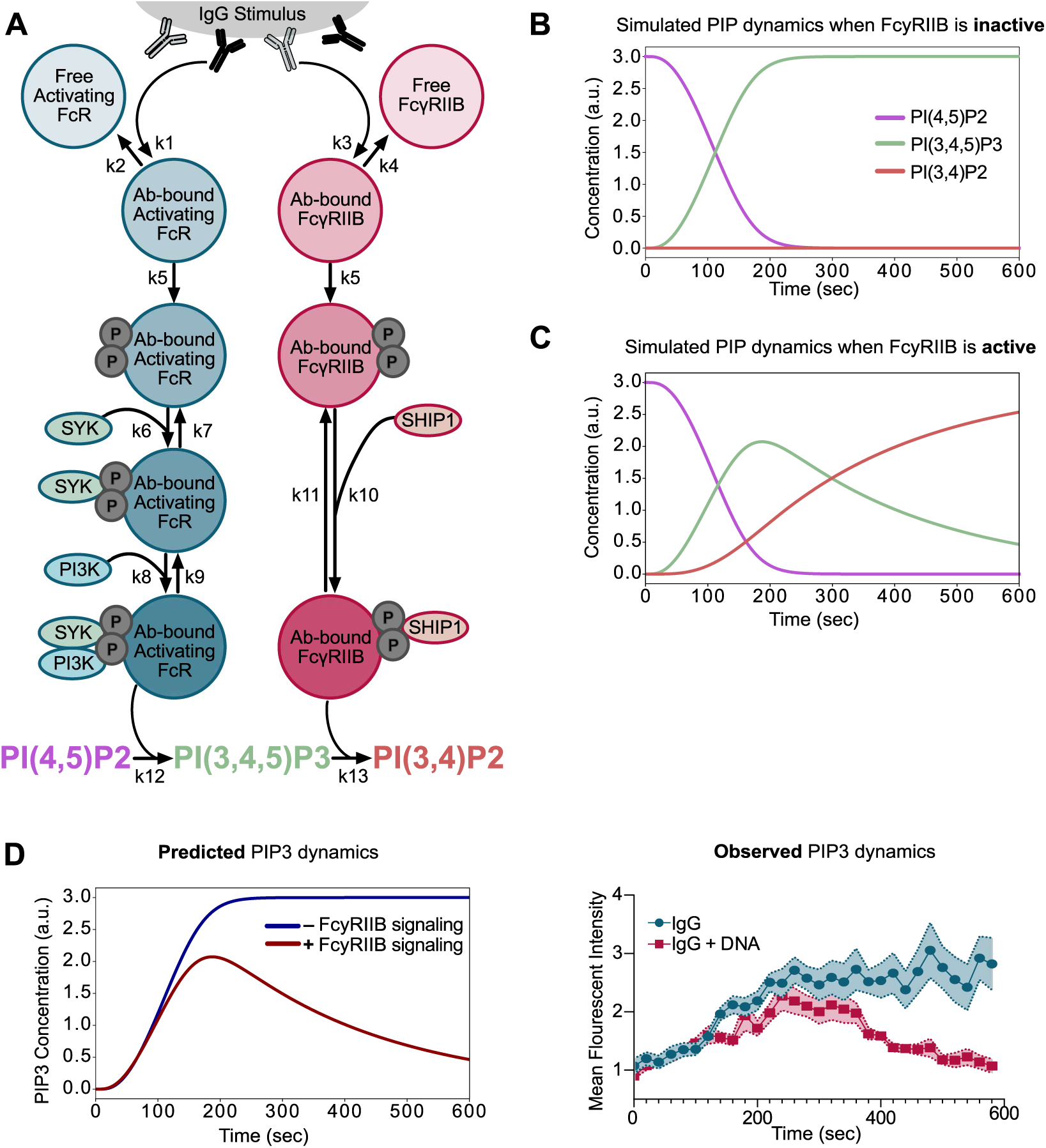
In Silico model of paired activating and inhibitory signaling predicts PIP3 dynamics at the phagocytic cup. **(A)** Schematic summarizes the in silico model. Rate constant values and definitions are indicated in Supplementary Tables 3 and 4. **(B-C)** Model output of PIP dynamics (PI(4,5)P2 in purple, PI(3,4,5)P3 in green, PI(3,4)P2 in red) with (C) and without (B) FcγRIIB input during a 600 second simulation of IgG stimulus. **(D)** Predicted (left) and observed (right) PI(3,4,5)P3 dynamics after IgG binding with (red) and without (blue) FcγRIIB activity. Observed values were determined by the intensity of the PIP3 fluorescent reporter (BTK-PH-GFP) as FcγRIIB DNA CAR macrophages encounter a bead containing IgG (blue) or IgG+DNA (red). Pairwise comparisons were determined by Mann Whitney U test. Each solid point represents the mean of 10 cells. Shaded regions represent mean +/- SEM.

To validate this prediction in cells, we measured PIP3 dynamics and fit them to our model predictions **(Fig. 2D, S5A-C)**. We fluorescently tagged the BTK PH domain, which specifically binds PIP3, and measured cup intensity over time during individual phagocytic events (*41*). PIP3 rapidly accumulates and then plateaus or slowly declines at phagocytic cups surrounding targets with only IgG opsonization. When FcγRIIB DNA CAR is active, PIP3 initially accumulates at a similar rate, but subsequently rapidly declines. This aligns with our in silico prediction that paired activating and inhibitory Fc Receptors generate a pulse of PIP3.

### Activating Fc Receptors control initiation of phagocytosis while inhibitory FcγRIIB controls the duration of signaling

If FcγRIIB simply dampened macrophage signaling, this could be accomplished more simply by expressing less activating Fc Receptors. In fact, macrophages alter Fc Receptor expression to adjust their appetite to many extracellular cues (*13*, *42*, *43*). To highlight this, we used publicly available databases to compare activating Fc Receptor and FcγRIIB expression across macrophage populations. Expression of both the activating and inhibitory Fc Receptors varied by more than an order of magnitude **(Table S6)** (*44*). Given this variability, we next asked whether activating and inhibitory Fc Receptors control distinct aspects of phagocytic signaling.

PIP3 must surpass a critical threshold to activate phagocytosis (*23*). Our model predicted that increasing inhibitory receptor signaling (phosphorylated FcγRIIB, **Table S3**) altered the duration of elevated PIP3 **(Fig 3A)**. This suggests that inhibitory signaling controls how long phagocytosis continues before it fails. To test this prediction in cells, we titrated FcγRIIB CAR DNA ligand on 9 µm diameter IgG beads, and measured the time macrophages engaged a target before phagocytosis failed. As DNA concentration increased, the time before phagocytosis failure decreased **(Fig 3B)**. Further, this shorter time corresponded to a lower percentage of the target surface area covered at failure.

**Figure 3.**
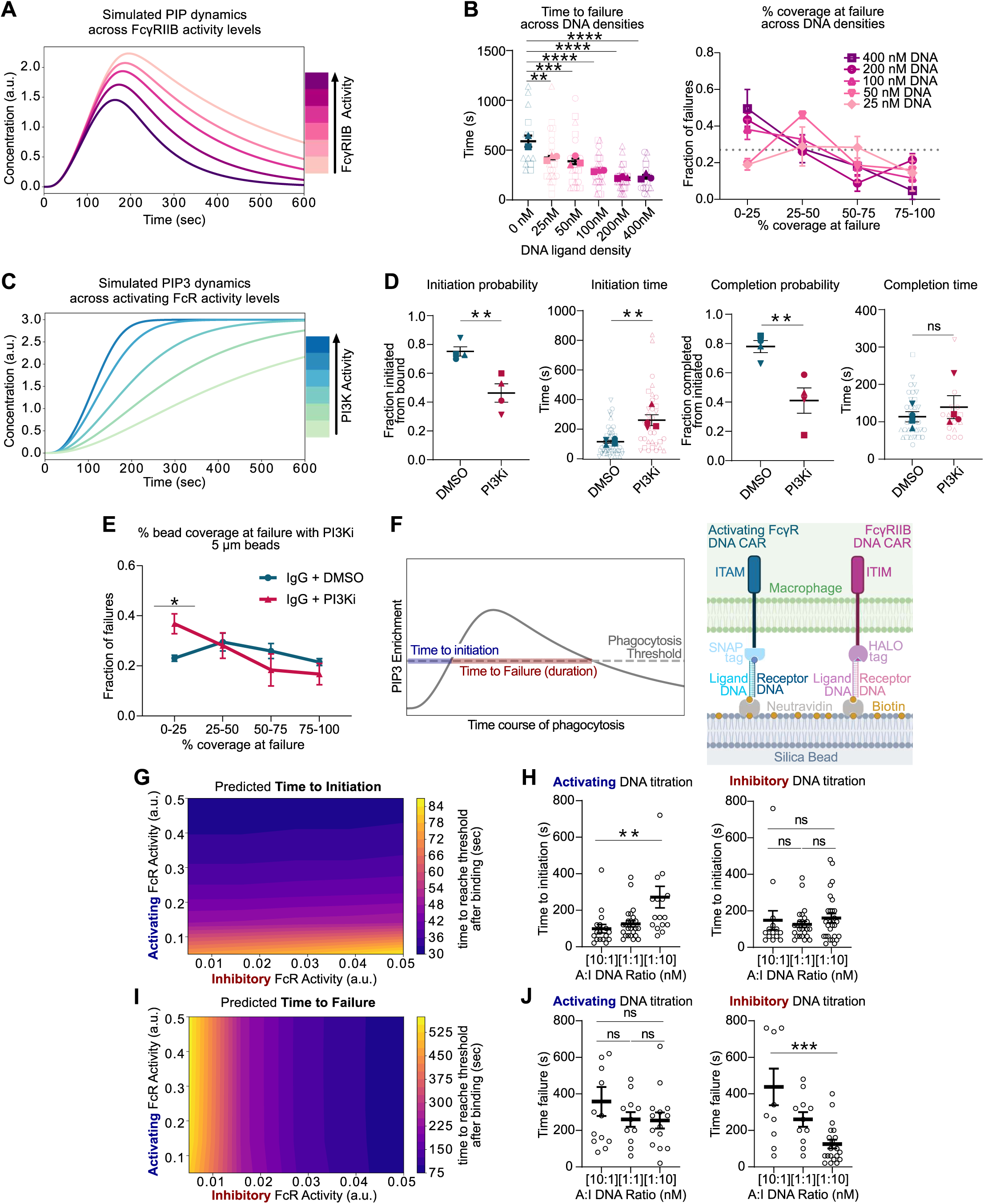
Activating and Inhibitory Fc Receptor signaling play distinct roles in coordinating phagocytosis. **(A)** Graph shows in silico prediction of PIP3 enrichment over time. Lines represent simulated PIP3 levels with increasing phosphorylated FcγRIIB. **(B)** FcγRIIB DNA CAR macrophages were incubated with 9 µm beads containing IgG and an increasing amount of FcγRIIB DNA CAR ligand. The time between initiation of phagocytosis and failure (left graph) and the bead coverage at failure (right graph) decreases with increasing ligand DNA as measured by timelapse microscopy. **(C)** Graph shows in silico prediction of PIP3 enrichment over time. Lines represent simulated PIP3 levels with increasing phosphorylated activating Fc Receptor. **(D)** After treatment with 50 µm LY294002, time lapse microscopy was used to quantify the fraction of beads bound to a macrophage that proceeded to initiation; the time from binding to initiation; the fraction of beads that proceeded from initiation to completion; the time between initiation and completion of phagocytosis. **(E)** The graph shows the target surface covered by the macrophage membrane at failure for IgG targets with (red) or without (blue) treatment with LY294002. **(F)** Schematics depict the parameters measured in G and I (left) and the DNA CAR system used in H and J (right). **(G)** Heatmap shows the predicted time to initiation for varying activating and inhibitory Fc Receptor inputs. **(H)** Supported lipid bilayer coated 9 µm diameter targets with the indicated ratios of activating and inhibitory DNA ligand were added to RAW264.7 cells expressing both the activating FcγR DNA CAR and FcγRIIB DNA CAR. The time between bead binding and initiating phagocytosis was monitored by timelapse microscopy. Data produced with the 1:1 ligand ratio is duplicated between graphs for comparison. **(I)** Heatmap shows the predicted time to failure of phagocytosis after initiation. **(J)** Activating and inhibitory DNA ligands were titrated as in (I). The time between initiating and failing phagocytosis was monitored by timelapse microscopy. Data produced with the 1:1 ligand ratio is duplicated between graphs for comparison. In B and E, points represent the mean of 3 independent experiments. In other graphs, solid data points represent the mean of an independent experiment, while unfilled data points represent individual cell measurements. Data points from replicates performed on the same day are denoted by symbol shape except in H and J. In all graphs, bars represent the mean +/- SEM. *indicates p<0.05, **indicates p<0.01, ***indicates p<0.001, ****indicates p<0.0001 by ordinary one-way ANOVA with Tukey’s multiple comparisons test (B, H, J), or unpaired two-tailed t test (D, E)

We then used an *in silico* model to vary the strength of the activating Fc Receptor input. Varying the activating Fc Receptor arm (phosphorylated activating FcγR, **Table S3**) primarily affected PIP3 generation speed **(Fig 3C)**. This suggests reducing activating Fc Receptors will slow or reduce initiation. To test this prediction, we slowed PIP3 generation using the PI3K inhibitor LY294002, using a concentration that inhibited phagocytosis to a similar extent as the FcγRIIB DNA CAR **(Fig S6A)**. As our model predicted, PI3K inhibition did not replicate the phenotype of FcγRIIB activation. Instead, PI3K inhibition significantly impacted both the probability and speed of initiating phagocytosis without impacting the morphology of the cup **(Fig 3D, S6B-C)**. If phagocytosis failed after initiation, failure most frequently occurred when the macrophage membrane covered less than 25% of the target **(Fig 3E)**. This suggests increasing FcγRIIB is phenotypically distinct from less Fc Receptor activation.

We then used our *in silico* model to predict PIP3 dynamics during phagocytosis across many different A:I signaling ratios, scanning across two orders of magnitude for each signaling arm. Because PIP3 levels must surpass a critical threshold to initiate phagocytosis (*23*), we set an arbitrary PIP3 threshold at half the maximum PIP3 level and asked how titrating activating and inhibitory inputs affected the time to reach this threshold as a proxy for time to initiate phagocytosis **(Fig 3F)**. Our in silico model predicted that titrating the activating signaling arm strongly impacted the time to initiate phagocytosis, while titrating the inhibitory signaling arm had a comparatively small effect (**Fig 3G)**.

To test this experimentally, we used our synthetic DNA CAR system **(Fig 3F)**. We generated RAW264.7 macrophages expressing the FcγRIIB DNA CAR and an activating Fc Receptor CAR with an extracellular SNAP tag paired to the transmembrane and intracellular ITAM domain of the common gamma chain (*16*). This activating CAR specifically binds a benzylguanine modified ssDNA, while the FcγRIIB DNA CAR binds a Halo Ligand modified ssDNA, each with a unique sequence **(Fig 3F)**. This system allowed us to separately titrate the complementary DNA ligands. Varying the activating ligand affected time to initiation while varying inhibitory ligand did not **(Fig 3H)**.

We then used our model to predict how long PIP3 remained above our arbitrary threshold as a proxy for the time before phagocytosis fails. Our model predicted that inhibitory signaling strongly impacts time to failure, while activating signaling had only a modest effect **(Fig 3I)**. We again tested this prediction using DNA CARs. Varying the inhibitory ligand affected time to failure, while varying the activating ligand did not **(Fig 3J)**. Overall, these data suggest changing activating Fc Receptor expression or IgG affinity primarily affects initiation of signaling, or macrophage sensitivity, while changing inhibitory FcγRIIB expression or engagement primarily affects signaling duration.

### FcγRIIB regulates the duration of ERK activation

FcγRIIB polymorphisms are associated with inflammatory disorders in humans (*5*, *7*). We hypothesized that an FcγRIIB-driven time limit may selectively regulate responses that require sustained Fc Receptor signaling, like inflammatory cytokine secretion, while allowing macrophages to continue phagocytosing bacteria-sized targets. To test this, we incubated FcγRIIB sgRNA and nontargeting sgRNA macrophages with 1.86 micron diameter beads, and measured phagocytosis and TNFα secretion **(Fig. 4A, S7A)**. As expected, FcγRIIB sgRNA did not measurably affect phagocytosis of these small targets. In contrast, FcγRIIB sgRNA macrophages increased TNFα production in response to IgG1, while control macrophages did not.

**Figure 4.**
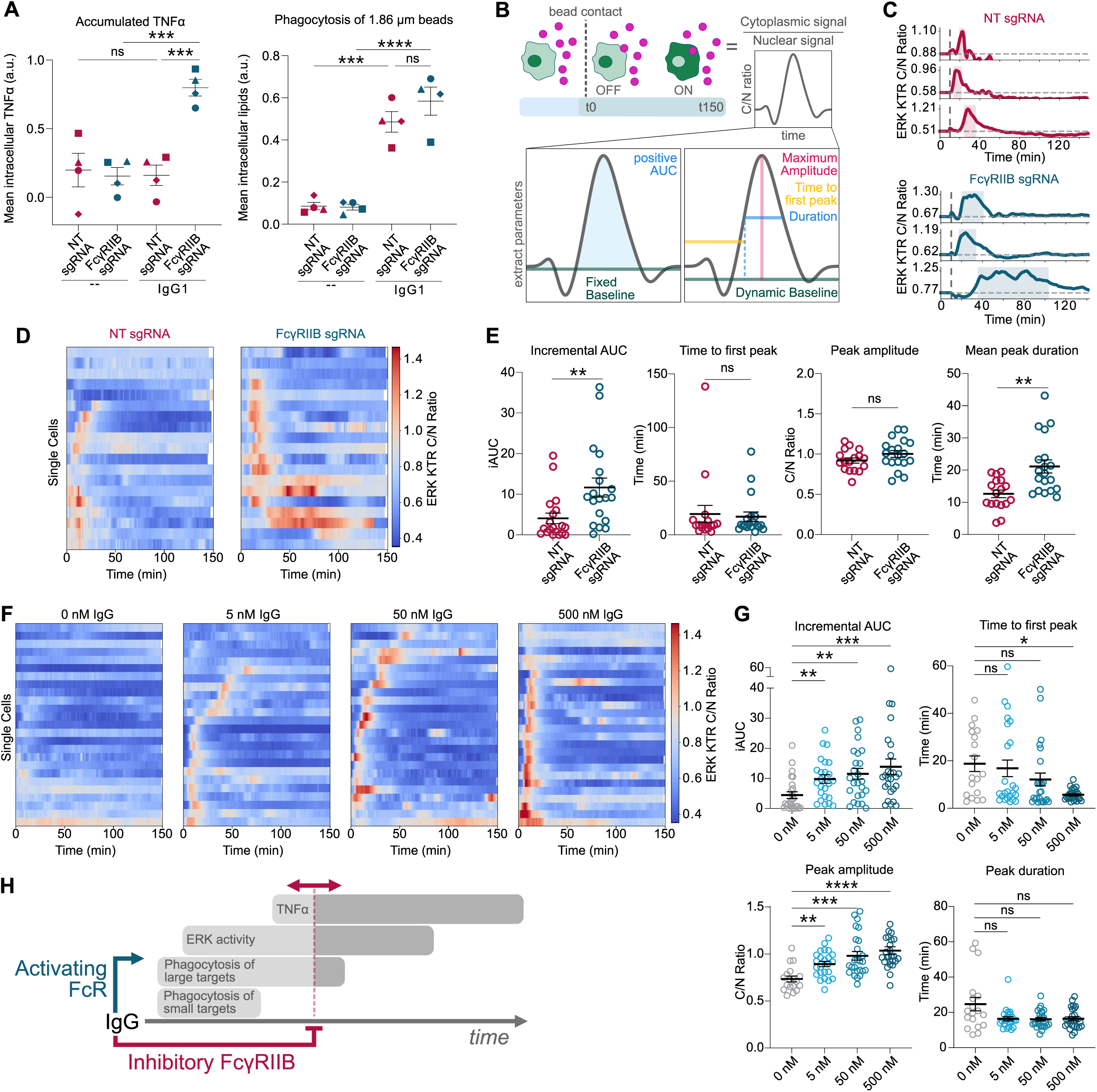
FcγRIIB regulates duration of the ERK response. **(A)** HoxB8 macrophages infected with FcγRIIB KO (blue) or NT sgRNA (red) were incubated with either un-opsonized or IgG1 1.86 µm diameter beads. Internalized lipid fluorescence was measured after 30 minutes as a proxy for phagocytosis (left). TNF⍺ was measured after an additional 4 hours of Brefeldin A treatment, followed by immunostaining for intracellular TNF⍺ accumulation. **(B)** Schematic of ERK analysis in (E,G). The cytoplasmic to nuclear (C/N) ratio of the ERK KTR sensor was measured by timelapse microscopy. The time to first peak, maximum amplitude, and duration was quantified from individual traces. **(C)** Representative traces show the C/N ratio of the ERK KTR sensor in HoxB8 macrophages infected with FcγRIIB or NT sgRNA after engulfing IgG1 beads. Traces represent individual cells. Vertical dashed lines indicate T0, when the macrophage contacts the IgG1 bead. **(D)** Heatmaps depict ERK KTR C/N ratio over time as described in (C). Each row is an individual cell. **(E)** Graphs display incremental area under the curve (AUC), time to first peak, peak amplitude, and peak duration for C/N ratio traces shown in D. **(F)** Heatmaps show C/N ratio of ERK-KTR-GFP WT HoxB8 macrophages phagocytosing beads with varying IgG1 density. For comparison, beads D,E in were conjugated to 500 nM IgG. **(G)** Graphs display incremental area under the curve (AUC), time to first peak, peak amplitude, and peak duration for C/N ratio traces shown in F. **(H)** Schematic summarizes finding that FcγRIIB limits the duration of IgG response, allowing phagocytosis of small targets to proceed while inhibiting phagocytosis of large targets and inflammatory cytokine secretion. In A, solid points represent the mean of independent experiments. Data points from replicates performed on the same day are denoted by symbol shape. For other graphs, each data point represents a different cell, collected across 3 independent experiments. In all graphs, bars represent the mean +/- SEM. *indicates p<0.05, **indicates p<0.01, ***indicates p<0.001 by ordinary one-way ANOVA with Tukey’s multiple comparisons test (A), Welch’s unpaired t test (E), or Kruskal-Wallis test with Dunn’s multiple comparisons test (G).

IgG immune complexes drive inflammation by activating the MAPK pathway downstream of the Fc Receptor (*45*). The MAPK pathway is highly sensitive to signaling dynamics, with ERK pulse amplitude and duration encoding distinct functional outcomes (*46–50*). To test if FcγRIIB impacts ERK kinetics, we used the ERK-KTR reporter, which localizes to the nucleus when ERK is inactive and the cytoplasm when ERK is active providing a rapid, sensitive read out of ERK activity over time in single cells **(Fig. 4B)**(*51*). Control macrophages phagocytosing IgG beads displayed a short pulse of ERK activation **(Fig. 4C,D)**. The same stimulus elicited a more robust ERK response in FcγRIIB sgRNA cells. To quantify differences in ERK kinetics, we extracted the area under the curve (AUC; total activity), the time from bead contact to the first ERK activity peak, the maximum ERK activation (amplitude), and the time ERK activity remained above baseline (duration) **(Fig. 4E, S7B-F)**. FcγRIIB sgRNA cells had increased total ERK activity (AUC). This increase was driven by an increase in ERK duration. The average peak amplitude and the time to first peak remained the same. Overall, this demonstrates that FcγRIIB regulates duration but not initiation of signaling, similar to the effect on phagocytosis.

We next asked if increasing Fc Receptor activation by increasing total IgG caused a similar change in ERK duration. Higher IgG density again increased total ERK activity **(Fig. 4F,G, S7G-H)**. However, increasing IgG did not affect the duration of ERK activity. Instead, high IgG increased peak amplitude and shortened the time to the first peak. This suggests that activating and inhibitory Fc Receptors have distinct effects on ERK kinetics.

## Discussion

The A:I ratio has informed antibody engineering since it was proposed two decades ago (*14*, *15*). Our data updates this model, suggesting an antibody’s effect is not simply the balance of activating and inhibitory Fc receptor engagement. Instead activating Fc Receptors control macrophage sensitivity while FcγRIIB controls signaling duration.

A FcγRIIB-driven time limit could impact macrophage biology in ways that a simple dampener would not. First, a time limit would terminate signaling if a target cannot be phagocytosed. This could free macrophages to later engage with a new target, and prevent endless cytokine secretion from a single target interaction. Second, FcγRIIB may selectively inhibit macrophage responses that require sustained signaling, allowing macrophages to fine-tune their response to IgG **(Fig. 4H)**. In our experiments, phagocytosis of small targets, like bacteria or red blood cells, was not affected by FcγRIIB. In contrast, phagocytosis of large targets and TNFα production are inhibited by FcγRIIB. Different ERK kinetics are associated with different transcriptional profiles in multiple immune populations, suggesting that altering ERK dynamics could change which genes IgG activates. Titrating FcγRIIB engagement or expression would allow macrophages to change the duration of signaling, adjusting their response based on the environment or a specific target.

FcγRIIB functions differently than SIRPA, another ITIM-containing receptor, even when suppressing the same activating signal. SIRPA alters phagocytosis speed and cup morphology, while FcγRIIB does not (*25*). Inhibitory receptors are frequently described as a straightforward counterbalance to activating receptors, but these findings argue for a more nuanced view where different inhibitory receptors inhibit distinct downstream nodes. This suggests therapeutics targeting inhibitory receptors should consider the target’s signaling mechanism, not just when and where it is expressed.

## Acknowledgments

We thank members of the Morrissey lab for thoughtful discussion. We thank Ben Winer (NYU) for HoxB8 macrophage progenitors. lenti-sgRNA blast was a gift from Brett Stringer. pLenti6-H2B-mCherry was a gift from Torsten Wittmann.

## Author Contributions

Conceptualization, A.B., E.T.S., and M.A.M.

Methodology, A.B., E.T.S., and M.A.M.

Software, A.B., E.T.S., A.M., M.Z.W., E.Y.

Investigation, A.B., E.T.S., C.H., K.K.

Visualization, A.B., E.T.S.

Funding acquisition, M.A.M., M.Z.W., E.Y.

Supervision, M.A.M.

Writing – original draft, A.B., E.T.S., M.A.M.

Writing – review and editing, A.B., E.T.S., A.M., K.K., M.Z.W., E.Y, and M.A.M.

## Data, code, and materials availability

All data are available in the main text or the supplementary materials. All reagents and materials are available from M.A.M. upon reasonable request. All code and associated data will be made publicly available upon final publication.

## Materials and Methods

### Cell lines

HEK293T cells specialized for Lentivirus production were obtained from Takara (Cat# 632180)\ and were cultured DMEM media containing 10% FBS, 1% PSG. RAW264.7 (ATCC TIB-71) cells were obtained from ATCC and were cultured in DMEM media containing 10% FBS, 1% PSG, 1mM sodium pyruvate. All cells were maintained at 37°C with 5% CO2. Cells were kept below 20 passages for experiments. HoxB8 immortalized myeloid progenitors were generously gifted from Benjamin Winer and were prepared as described in Settle et al. (Settle et al., 2024) according to an established protocol (Wang et al., 2006). In short, ER-HoxB8 progenitor cells were generated from bone marrow harvested from ROSA-Cas9 mice (Jackson Labs #026179, 6– 8 weeks old) and retrovirally transduced with the ER-HoxB8 oncogene. Cells were then cultured in RPMI supplemented with 1% pen–strep, 2 mM l-glutamine, 10% FBS, 1 µM β-estradiol, 5% GM-CSF conditioned media for 1 month to isolate immortalized clones expressing HoxB8-ER.

### FcγRIIB KO and purification

To generate CRISPR KO lines, HoxB8 Cas9 cells were infected with lentivirus containing targeting (KO) or non-targeting (NT) sgRNA and grown in HoxB8 culture media. After 72 h, the media was replaced with fresh HoxB8 Culture Media with 5 µg/mL blasticidin-HCl (Thermo Fisher A1113903) for selection of FcγRIIB KO or NT cells. After selection and differentiation, we measured FcγRIIB expression by flow cytometry using the Anti-CD32b APC antibody (Invitrogen Cat# 17-0321-80) and APC isotype control (Invitrogen Cat# 17-4724-81). We also verified this by qPCR, collecting 1 million FcγRIIB KO, NT and WT cells per replicate. RNA extraction was done using Qiagen RNeasy kit Cat# 74104 and cDNA synthesis was done using BioRad iScript cDNA Synthesis Kit Cat# 1708890. qPCR reactions were performed using SsoAdvanced Universal SYBR Green Supermix (BioRad Cat#1725270) in a 10 uL total reaction volume. qPCR assay was done using IDT PrimeTime predesigned qPCR Primer Assays at a final concentration of 500 nM for all variants of mouse *Fcgr2b* (Mm.PT.58.8598996) and *Gapdh* housekeeping gene (Mm.PT.39a.1) (see Oligos section). The thermocycler was set for 1 cycle of 95℃ for 3 minutes, and 40 cycles of 95℃ for 10 seconds and 60℃ for 20 seconds. qPCR results were analyzed using the 2^(-ΔΔCq) method (Livak and Schmittgen, 2001).

### Lentivirus production and infection

All constructs were expressed in RAW264.7 macrophages using lentiviral infection. Lentivirus was produced in HEK293T cells transfected with pMD2.G (Gift from Didier Trono, Addgene plasmid # 12259 containing the VSV-G envelope protein), pCMV-dR8.258 (Gift from Bob Weinberg, Addgene plasmid #8455), and a lentiviral backbone vector containing the construct of interest using lipofectamine LTX (Invitrogen, Cat# 15338–100). The media was harvested 72 h post-transfection, filtered through a 0.45 µm filter (Millapore, Cat# SLHVM33RS) and concentrated using LentiX (Takara Biosciences, Cat# 631232). Concentrated lentivirus was added to cells on day 4 of differentiation in HoxB8 cells.

### Supported lipid bilayer coated beads

#### SUV preparation

For anti-biotin IgG conjugated beads the following chloroform-suspended lipids were mixed and desiccated overnight to remove chloroform: 97.8% POPC (Avanti, Cat# 850457), 2% biotinyl cap PE (Avanti, Cat# 870273), 0.1% PEG5000-PE (Avanti, Cat# 880230, and 0.1% atto390- DOPE (ATTO-TEC GmbH, Cat# AD 390–161) or 0.1% atto647-DOPE (ATTO-TEC GmbH, Cat# AD 647–161). For anti-CD20 conjugated beads: 97.8% POPC (Avanti, Cat# 850457), 2% DOGS-NTA (Avanti, Cat# A89404), 0.1% PEG5000-PE (Avanti, Cat# 880230, and 0.1% atto647-DOPE (ATTO-TEC GmbH, Cat# AD 647–161). The lipid sheets were resuspended in PBS, pH7.2 (GIBCO, Cat# 20012050) at 10 mM concentration and stored under inert nitrogen gas. For all SUVs, the lipids were broken into small unilamellar vesicles via several rounds of freeze-thaws. The lipids were then stored at −80°C under nitrogen gas. To remove aggregated lipids, the solution was diluted to 2 mM and filtered through a 0.22 µm filter (Millapore, Cat# SLLG013SL) immediately prior to use.

#### Bead preparation

Silica beads with a 5.01 µm diameter (10% solids, Bangs Labs, Cat# SS05003, Lot # 16595) or 9.2 µm diameter (Cospheric, Cat# SiO2MS-2.0 9.2-1g) were washed several times with PBS, mixed with 1mM SUVs in PBS and incubated at room temperature for 30 min with end-over-end mixing to allow for bilayer formation. Beads were then washed with PBS to remove excess SUVs and incubated in 0.2% casein (Sigma, Cat# C5890) in PBS for 15 min before protein coupling (Neutravidin or human CD20, anti biotin IgG1 or anti hCD20 IgG1, IgG2a). When indicated, beads with various diameters were used (1.86um diameter, Cospheric, Cat# SiO2MS-2.0 1.86-1g; 4.3um diameter, Cospheric, Cat# SiO2MS-2.0 4.3-1g; 7.75um diameter, Cospheric, Cat# SiO2MS-2.0 7.75-1g; 9.2um diameter, Cospheric, Cat# SiO2MS-2.0 9.2-1g).

For IgG conjugated beads, we then added anti-biotin AlexaFluor647-IgG (Jackson ImmunoResearch Laboratories Cat# 200-602-211, Lot# 167722), antibiotin AlexaFlour488-IgG (Jackson ImmunoResearch Laboratories Cat#200-542-211, Lot# 174725), or anti-biotin IgG (unlabeled, Jackson ImmunoResearch Laboratories Cat# 200-002-211, Lot # 155583) at 500 nM to a 10x dilution of beads (1% solids), unless otherwise indicated. The number of IgG molecules per bead was previously estimated to be 10–18 IgG molecules/μm^2^ (Bond et al., 2023). For DNA conjugated beads, we then added 1 ug/ml neutravidin to beads after the addition of IgG followed by 250 nM biotinylated ligand DNA strand in 20 mM MgCl2 unless otherwise noted in figure legends. Between the addition of each conjugate, beads were washed 3x to remove unbound conjugates. For CD20 conjugated beads with various isotypes of IgG, 6xHis-tagged human CD20 (Genscript Cat# Z03630-100) was added at 500 nM to beads covered in a supported lipid bilayer containing 2% DGS NTA lipid. Beads were washed 3x in PBS to remove unbound CD20, then mouse anti-hCD20 IgG isotypes were added (Invivogen Cat# hcd20-mab9, hcd20-mab10) at a concentration of 500 nM unless otherwise noted. Equal IgG loading between isotypes was determined via flow cytometry using an anti-mouse IgG F(ab’)2 Fragment 647 Conjugate (Cell Signaling Cat# 4410).

### Phagocytosis assay

Approximately 80,000 RAW264.7 or 50,000 HoxB8 macrophages were plated in one well of a 96-well glass-bottom MatriPlate (Brooks, Cat# MGB096-1-2-LG-L) approximately 24 h before the start of the experiment. ∼8×10^5^ beads were added to wells and engulfment was allowed to proceed for 30 min. The cells were imaged using spinning disk microscopy (40 x 0.95 NA Plan Apo air). Internalized particles were identified by their fluorescent supported lipid bilayer, and counted in ImageJ by a blinded analyzer using Blind-Analysis-Tools-1.0 ImageJ plug-in. For all phagocytosis assays, independent experiments consist of 50-150 cells scored. Where indicated, phagocytosis was normalized such that each point represents the mean of 2 technical replicates, normalized to the highest technical replicate that day (n=50-150 cells per technical replicate).

### DNA receptor strand synthesis and conjugation

Receptor strands were synthesized and conjugated to DNA CARs as previously described (Farlow et al., 2013; Kern et al., 2021). Briefly, the ligand strand oligonucleotides for both the activating FcR DNA CAR (SNAP linker), and the inhibitory FcγRIIB DNA CAR (HALO linker) were ordered from IDT conjugated with 5’ terminal amine (5amMC6) and diluted in 0.15 M HEPES pH 8.5 to a final concentration of 2 mM. The 5’ amine of the FcR DNA CAR receptor strand was conjugated to N-hydroxysuccinimide ester (BG-GLA-NHS)-functionalized benzylguanine at a 1:50 oligo:BG-GLA-NHS ratio overnight, then purified in an illustra NAP-5 Columns (Cytiva, Cat# 17085301), using H2O for elution. The 5’ amine of the FcγRIIB DNA CAR receptor strand was conjugated the same, except using a HaloTag Succinimidyl Ester Ligand (Promega Cat# p1691) in place of BG-GLA-NHS. Prepared receptor strands were added to cells at 1 µm in serum-free DMEM for 10 min before being washed out.

### Inhibitors

For SHIP1/2 inhibition, 3AC (Sigma, Cat# 565835, 10 µM) and AS1938909 (Sigma, Cat#565840, 5 µM); for SHP1/2 inhibition, TPI-1 (Sigma, Cat# SML3119, 4 µM) and SHP099 (MedChemExpress, cat# HY-100388, 7 µM) were added to cells 24 hours prior to the start of the experiment. For Dynamin inhibition, MiTMAB (Sigma, cat#324411, 1 µm) was added to cells 15 min prior to the start of the experiment. For PI3K inhibition, LY294002 (Sigma, cat#440202, 50 µM) was added to cells 30 min prior to the start of the experiment.

### Live imaging assays

#### Timelapse imaging of phagocytosis kinetics and percent coverage at failure

RAW264.7 or HoxB8 macrophages were plated as for a phagocytosis assay. Individual positions were selected manually using ND acquisition in Elements before beads were added. Phagocytosis was imaged at 20s, 1m, or 5m intervals (for kinetics, failure time, percent coverage respectively) through at least 10 z planes for 15-30 min. Only beads that bound with more than 5 min remaining were included in each data set. Binding was defined as the first frame at which the bead surface contacts macrophage membrane, initiation was defined as the first frame at which the macrophage membrane extends around the target, completion was defined as the first frame where macrophage membrane completely covers the target. Failure was defined as the first frame at which retraction or indefinite stalling of the cup was observed. For percent coverage analysis, timelapse hyperstacks were compressed into maximum projections in z, to view all cup projections in each frame for visually determining the % coverage around the bead. For all kinetics and percent coverage analysis, independent experiments consist of 10-50 bead contacts.

#### PI(3,4,5)P3 Localization

Approximately 80,000 RAW264.7 macrophages were plated in one well of a 96-well glass bottom MatriPlate (Brooks, Cat# MGB096-1-2-LG-L) between 12 and 24 hours prior to the experiment. 8×10^5^ 5 µm silica beads were added to wells and cells. Z-stacks were acquired every 20 seconds over a 30-minute time-lapse immediately following the addition of beads using a 100 x 1.49 NA oil immersion objective. PI(3,4,5)P3 enrichment was determined by measuring the mean intensity macrophage membrane contacting the bead contacts normalized to the adjacent membrane using the freehand line analyzer tool in ImageJ. Enrichment values (mean grey values) were acquired at each time point, and background subtracted, and used to calculate the fold change enrichment at the cup.

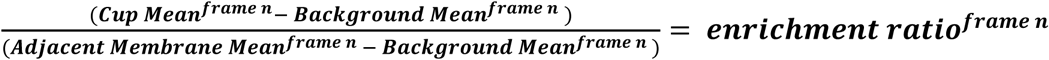

### Assessing TNFαlpha Production

#### TNFαlpha bead assay and staining protocol

Approximately 50,000 HoxB8 macrophages were plated per well in 96-well glass-bottom MatriPlates (Brooks, Cat# MGB096-1-2-LG-L) 24 h before the start of the experiment. For each experiment, two sets of wells were prepared in parallel: one for intracellular TNFα staining and one for scoring bead engulfment, the latter necessary because detergents used in antibody staining dissolve the supported lipid bilayer and renders the beads no longer visible in the stained samples. Macrophages in the TNFα-staining wells were loaded with CellTracker Orange CMTMR (ThermoFisher Cat# C2927) and those in the engulfment wells with CellTracker Green CMFDA (ThermoFisher Cat# C2925), in each case by aspirating the media and replacing it with 1 µM dye in warm, serum-free RPMI for 10 min at 37 °C, then washing into complete media.

Approximately 8×10⁵ unopsonized or IgG1 opsonized 1.86 µm diameter supported lipid bilayer coated beads were added per well and phagocytosis was allowed to proceed for 30 min at 37 °C. The engulfment wells were imaged live immediately after this 30 min incubation by spinning disk confocal microscopy. In the TNFα-staining wells, Brefeldin A (ThermoFisher Cat# B5936) was then added directly to the wells without washing out beads to a final concentration of 7.5 µg/mL and incubated for 4 h at 37 °C to trap intracellular cytokines. Cells were fixed with 4% cold paraformaldehyde in PBS for 15 min, permeabilized with 0.1% Triton X-100 in PBS for 15 min, and blocked in blocking buffer (0.1% Triton X-100, 1% BSA in PBS) for 15 min, all at room temperature. Cells were then incubated with an AlexaFluor488-conjugated anti-mouse TNFα antibody (ThermoFisher Cat# 53-7321-82) diluted 1:500 in a blocking buffer overnight at 4 °C, protected from light. A matched AlexaFluor488 rat IgG1κ isotype control (BioLegend, Cat# 400417) was stained in parallel. The following day, cells were washed 3× in PBS, counterstained with 300 nM DAPI in PBS for 5 min at room temperature, and washed 3× in PBS. Cells were imaged by spinning disk confocal microscopy and per-cell TNFα fluorescence intensities were quantified as described below. For each technical replicate, 20 fields of view were scored per condition.

#### TNFα production analysis

Macrophages were identified in TIFF images using a semi-automated pipeline built on ilastik v1.4.2, developed with a custom Python package, and structurally inspired by the nucleus-seeded region-growing strategy used to identify whole-cell macrophage boundaries (Joffe et al., 2020). Nuclei were segmented from the nuclear channel by Gaussian smoothing followed by Otsu thresholding, producing a binary nuclear mask. Independently, each field of view was classified using an ilastik Pixel Classification model trained on manually annotated cytoplasmic-channel images, producing a continuous membrane probability map. These two outputs were then combined: the nuclei served as seed points, and the membrane probability map served as the surface for a marker-controlled watershed algorithm, which grew each cell outward from its nucleus until it reached the cell boundary indicated by the membrane probability. Watershed growth was constrained to a foreground region combining thresholded membrane probability with the nuclear mask, so segmentation stayed within areas supported by real membrane or nuclear signal. Fields of view with an unusually high nucleus count or membrane area fraction were flagged and excluded from downstream analysis. Segmentation quality was confirmed by visual inspection of the 540 channel alongside detected nuclei and final cell boundaries for a subset of fields. For each identified cell, mean fluorescence intensity was measured across the full segmented cell area in the Green-488 channel. For each replicate 20 fields of view were scored per condition. To remove background, the mean 488 intensity of isotype control was subtracted. The mean for each condition was normalized to the max technical replicate in that experiment.

#### Engulfment/internalized lipid analysis

Macrophages were identified from TIFF images using a semi-automated pixel classification and instance segmentation pipeline using a custom Python package built on ilastik v1.4.1. Each field of view was then classified into cytoplasm, background and cell-contact classes using ilastik Pixel Classification work flow trained on independently annotated images of the cytoplasmic channel to generate a binary foreground mask. Segmentation quality was confirmed by visual inspection of quality-control overlays showing the input image and final cell instance boundaries for a representative subset of fields. For each point within a well, mean fluorescence intensity (MFI) of the well was calculated by measuring the total fluorescence intensity in the Far Red-640 channel of the cell segmented area divided by the area segmented as cell membrane. To correct for inter-replicate differences in absolute signal, each field’s background-subtracted MFI was divided by the average MFI of that replicate’s 5 brightest fields (pooled across all conditions), giving a normalized MFI relative to that replicate’s own high-signal reference.

### Assessing ERK activity

#### ERK activity Analysis

Approximately 50,000 HoxB8 macrophages expressing a kinase translocation reporter for ERK activity (addgene #59150) and an mCherry H2B nuclear marker (addgene #89766) were plated in one well of a 96-well glass bottom MatriPlate (Brooks, Cat# MGB096-1-2-LG-L) between 12 and 24 hours prior to the experiment. 30 minutes prior to the experiment media was replaced with serum and phenol free RPMI (Gibco cat # 11-835-030). 8×10^5^ 5 µm silica beads with IgG, unopsonized beads, or unopsonized beads and 10 ng/mL LPS (Sigma cat #L4516) was added to wells and cells were imaged using spinning disk confocal microscopy. Images were acquired every 1 minute over a 2.5 hour timelapse.

TIFF time-lapse videos were imported into Napari (v0.0.6), where a custom Python script segmented nuclei from the H2B channel and defined a cytoplasmic ring around each nucleus. We analyzed cells that expressed both reporters, showed minimal baseline ERK activity, and phagocytosed at least one bead within the first 10 frames. This selection was applied uniformly across all conditions, including unopsonized (blank) beads, where phagocytosing cells represent rare outliers within a largely non-phagocytic population. For every frame, mean ERK-KTR intensity was measured in the nucleus and cytoplasm, and the cytoplasm-to-nucleus (C/N) ratio was calculated as a readout of ERK activity. Segmented nuclei were linked across consecutive frames into single-cell trajectories using trackpy (v0.7) to produce single-cell C/N traces. Traces were then exported and analyzed with a separate Python script (Python 3.12.3). Each trace was smoothed, and four features were extracted using SciPy (v1.16.3): integrated area under the curve (overall ERK activity above baseline), peak amplitude, peak duration (full width at half maximum; FWHM), and time to first peak. Because integrated AUC reflects cumulative activity over the full trace, it was referenced to a single fixed baseline (mean of the first three timepoints), by convention, whereas the peak-based features (amplitude, duration, time to first peak) were referenced to each peak’s local half-maximum. Peaks were detected with SciPy’s “find_peaks” using a prominence threshold and a minimum spacing of 3 frames between peaks, and peak amplitude, duration, and timing were then measured from the full width at half maximum (50% relative height) of each detected peak, similarly to other studies analyzing ERK-KTR dynamics (Goglia et al., 2020). NumPy (v2.0.2), pandas (v2.2.2), and Matplotlib (v3.10.0) were used for numerical processing and figure generation.

### In Silico Modeling of Phospholipid Dynamics

#### Core Model Components

We modeled phospholipid dynamics with a Python-based ODE system, building on Suter et al.’s in silico approach to FcR signaling and phagocytosis (Suter et al., 2021). The core model emulates the dual signaling cascade triggered by IgG binding and integrates an activating and inhibitory signaling arm that converges on PIP3. **Table S3** describes the dynamics of receptor-antibody binding and phosphorylation of ITIMs and ITAMs, (Li et al., 2007; Maenaka et al., 2001; Barua et al., 2012); SYK, PI3K and SHIP1 activation (Barua et al., 2012; Faeder et al., 2003; Zhang et al., 2009; Panayotou et al., 1993; Ladbury et al., 1995), and rates of PIP3 generation and depletion (Waddell et al., 2023; Maheshwari et al., 2017; Huang et al., 2011). Our model simulates that phagocytic cup, which is separated from the rest of the plasma membrane by a diffusion barrier (Golebiewska et al., 2011). We did not include PTEN or PIP5K, the 3-phosphatase that would recycle PIP3 back to PI(4,5)P2 and the kinase that synthesizes PI(4,5)P2, since they are not localized to the cup during phagocytosis (Kamen et al., 2007; Fairn et al., 2009). PLCγ was excluded as a PIP2 sink, as it does not have a significant effect on phagocytosis (Cheeseman et al., 2006). In our model, the local PIP2 pool would need to be replenished before the same patch of membrane could support another round of phagocytic PIP3 signaling (Szymańska et al., 2008), which would likely happen when a target is released and the phagocytic cup equilibrates with the surrounding plasma membrane.

Parameter values assigned from published binding and enzymatic rate constants are described in **Table S4**. In Fig. 3A simulated FcγRIIB activity was titrated by altering the amount of phosphorylated FcγRIIB (FcRp_B in model code; phosphorylated ‘’Ab-bound FcγRIIB” in Fig. 2A schematic). Similarly, in Fig. 3C, simulated activating FcR activity was titrated by altering the amount of phosphorylated activating FcRIIIA (FcRp_A in model code; phosphorylated ‘’Ab-bound Act. FcR” in Fig. 2A schematic)

#### Single cell trace fitting

The model was fit to single-cell PIP3 intensity traces at the phagocytic cup (n = 10 cells per condition, 30 time points, 600 s). Raw traces were Savitzky-Golay smoothed and baseline-subtracted per cell. All 15 rate constants were freed per cell, initialized at previously reported values (**Table S4**) and constrained by non-negativity bounds. Traces were fit by nonlinear least squares (scipy.optimize.least_squares). Integration used solve_ivp with the implicit Radau method to handle stiffness.

#### Identifiability and condition comparison

Because all 15 rate constants were freed, individual parameters were not separately identifiable from a single PIP3 observable. To determine which parameter combinations the data constrain, a Fisher information matrix (FIM) was constructed in log-parameter space with the literature-reported values as a shared anchor for both conditions. The matrix was eigendecomposed, and its eigenvalues spanned many orders of magnitude; “stiff” (well-determined) directions were separated from “sloppy” (poorly-determined) directions at the largest gap in the log-eigenvalue spectrum to define an identifiable subspace. Condition-dependent differences were assessed on biologically interpretable parameter combinations (ratios and products of rate constants; such as a given enzyme’s recruitment), each represented as a direction in log-parameter space and projected onto the stiff subspace. Only combinations that were ≥75% within the subspace were reported. For each cell, a combination’s value was computed from that cell’s own fitted rate constants, expressed as log deviations from the anchor and projected onto the combination direction. These single cell values were then compared between conditions.

All computations used Python 3.12, SciPy 1.16, NumPy 2.0, and Matplotlib 3.10.

### Microscopy

Images were acquired on a spinning disk confocal microscope (Nikon Ti2-E inverted microscope with a Yokogawa CSU-W1 spinning disk unit and an Orca Fusion BT scMos camera) with a 40 x 0.95 NA Plan Apo air or a 100 x 1.49 NA oil immersion objective. The microscope has a stage top incubator for temperature, CO2 and humidity control from OkoLabs and a piezo Z drive. Images were acquired using Nikon NIS Elements.

### Quantification and statistical analysis

Statistical analysis was performed in Prism 8 (GraphPad). The statistical test used is indicated in the relevant figure legend. Sample sizes were predetermined and indicated in the relevant figure legend.

## References

1. S. Gordon, Phagocytosis: An Immunobiologic Process. Immunity 44, 463–475 (2016).

2. K. Weiskopf, I. L. Weissman, Macrophages are critical effectors of antibody therapies for cancer. mAbs 7, 303–310 (2015).

3. J. C. Osorio, P. Smith, D. A. Knorr, J. V. Ravetch, The antitumor activities of anti-CD47 antibodies require Fc-FcγR interactions. Cancer Cell 41, 2051–2065.e6 (2023).

4. R. Dahan, E. Sega, J. Engelhardt, M. Selby, A. J. Korman, J. V. Ravetch, FcγRs Modulate the Anti-tumor Activity of Antibodies Targeting the PD-1/PD-L1 Axis. Cancer Cell 28, 285–295 (2015).

5. F. Nimmerjahn, J. V. Ravetch, Fcγ receptors as regulators of immune responses. Nat Rev Immunol 8, 34–47 (2008).

6. M. Daëron, O. Malbec, S. Latour, M. Arock, W. H. Fridman, Regulation of high-affinity IgE receptor-mediated mast cell activation by murine low-affinity IgG receptors. J Clin Invest 95, 577–585 (1995).

7. K. G. C. Smith, M. R. Clatworthy, FcγRIIB in autoimmunity and infection: evolutionary and therapeutic implications. Nat Rev Immunol 10, 328–343 (2010).

8. F. Li, P. Smith, J. V. Ravetch, Inhibitory Fcγ Receptor Is Required for the Maintenance of Tolerance through Distinct Mechanisms. The Journal of Immunology 192, 3021–3028 (2014).

9. P. E. H. Sharp, J. Martin-Ramirez, S. M. Mangsbo, P. Boross, C. D. Pusey, I. P. Touw, H. T. Cook, J. S. Verbeek, R. M. Tarzi, FcγRIIb on Myeloid Cells and Intrinsic Renal Cells Rather than B Cells Protects from Nephrotoxic Nephritis. The Journal of Immunology 190, 340–348 (2013).

10. A. S. Yilmaz-Elis, J. M. Ramirez, P. Asmawidjaja, J. Van Der Kaa, A.-M. Mus, M. D. Brem, J. W. C. Claassens, C. Breukel, C. Brouwers, S. M. Mangsbo, P. Boross, E. Lubberts, J. S. Verbeek, FcγRIIb on Myeloid Cells Rather than on B Cells Protects from Collagen-Induced Arthritis. The Journal of Immunology 192, 5540–5547 (2014).

11. R. Clynes, J. S. Maizes, R. Guinamard, M. Ono, T. Takai, J. V. Ravetch, Modulation of Immune Complex–induced Inflammation In Vivo by the Coordinate Expression of Activation and Inhibitory Fc Receptors. The Journal of Experimental Medicine 189, 179– 186 (1999).

12. M. Rumpret, J. Drylewicz, L. J. E. Ackermans, J. A. M. Borghans, R. Medzhitov, L. Meyaard, Functional categories of immune inhibitory receptors. Nat Rev Immunol 20, 771– 780 (2020).

13. A. Bond, M. A. Morrissey, Biochemical and biophysical mechanisms macrophages use to tune phagocytic appetite. Journal of Cell Science 138, JCS263513 (2025).

14. F. Nimmerjahn, J. V. Ravetch, Divergent Immunoglobulin G Subclass Activity Through Selective Fc Receptor Binding. Science 310, 1510–1512 (2005).

15. D. J. DiLillo, J. V. Ravetch, Fc-Receptor Interactions Regulate Both Cytotoxic and Immunomodulatory Therapeutic Antibody Effector Functions. Cancer Immunol Res 3, 704–713 (2015).

16. N. Kern, R. Dong, S. M. Douglas, R. D. Vale, M. A. Morrissey, Tight nanoscale clustering of Fcγ receptors using DNA origami promotes phagocytosis, eLife (2021). 10.7554/eLife.68311.

17. R. Dong, T. Aksel, W. Chan, R. N. Germain, R. D. Vale, S. M. Douglas, DNA origami patterning of synthetic T cell receptors reveals spatial control of the sensitivity and kinetics of signal activation. Proc. Natl. Acad. Sci. U.S.A. 118, e2109057118 (2021).

18. M. J. Taylor, K. Husain, Z. J. Gartner, S. Mayor, R. D. Vale, A DNA-Based T Cell Receptor Reveals a Role for Receptor Clustering in Ligand Discrimination. Cell 169, 108–119.e20 (2017).

19. M. Ono, H. Okada, S. Bolland, S. Yanagi, T. Kurosaki, J. V. Ravetch, Deletion of SHIP or SHP-1 Reveals Two Distinct Pathways for Inhibitory Signaling. Cell 90, 293–301 (1997).

20. M. Ono, S. Bolland, P. Tempst, J. V. Ravetch, Role of the inositol phosphatase SHIP in negative regulation of the immune system by the receptor FeγRIIB. Nature 383, 263–266 (1996).

21. E. Muraille, P. Bruhns, X. Pesesse, M. Daëron, C. Erneux, The SH2 domain containing inositol 5-phosphatase SHIP2 associates to the immunoreceptor tyrosine-based inhibition motif of Fc gammaRIIB in B cells under negative signaling. Immunol Lett 72, 7–15 (2000).

22. S. Freeman, S. Grinstein, Promoters and Antagonists of Phagocytosis: A Plastic and Tunable Response. Annual Review of Cell and Developmental Biology 37, 89–114 (2021).

23. Y. Zhang, A. D. Hoppe, J. A. Swanson, Coordination of Fc receptor signaling regulates cellular commitment to phagocytosis. Proc. Natl. Acad. Sci. U.S.A. 107, 19332–19337 (2010).

24. J. A. Swanson, A. D. Hoppe, The coordination of signaling during Fc receptor-mediated phagocytosis. Journal of Leukocyte Biology 76, 1093–1103 (2004).

25. W. D. Miller, A. K. Mishra, C. J. Sheedy, A. Bond, B. M. Gardner, D. J. Montell, M. A. Morrissey, CD47 prevents Rac-mediated phagocytosis through Vav1 dephosphorylation. bioRxiv [Preprint] (2025). 10.1101/2025.02.11.637707.

26. P. Bruhns, B. Iannascoli, P. England, D. A. Mancardi, N. Fernandez, S. Jorieux, M. Daëron, Specificity and affinity of human Fcγ receptors and their polymorphic variants for human IgG subclasses. Blood 113, 3716–3725 (2009).

27. A. Mularski, R. Wimmer, F. Arbaretaz, G. L. Goff, M. Depierre, F. Niedergang, Dynamin-2 controls actin remodeling for efficient complement receptor 3-mediated phagocytosis. Biol Cell 115, e2300001 (2023).

28. F. Marie-Anaïs, J. Mazzolini, F. Herit, F. Niedergang, Dynamin-Actin Cross Talk Contributes to Phagosome Formation and Closure. Traffic 17, 487–499 (2016).

29. D. Kim, Y. Kwon, K. Cho, The biphasic behavior of incoherent feed-forward loops in biomolecular regulatory networks. BioEssays 30, 1204–1211 (2008).

30. S. Mangan, U. Alon, Structure and function of the feed-forward loop network motif. Proc. Natl. Acad. Sci. U.S.A. 100, 11980–11985 (2003).

31. U. Alon, Network motifs: theory and experimental approaches. Nat Rev Genet 8, 450–461 (2007).

32. D. Paul, S. Achouri, Y.-Z. Yoon, J. Herre, C. E. Bryant, P. Cicuta, Phagocytosis Dynamics Depends on Target Shape. Biophysical Journal 105, 1143–1150 (2013).

33. U. Golebiewska, J. G. Kay, T. Masters, S. Grinstein, W. Im, R. W. Pastor, S. Scarlata, S. McLaughlin, Evidence for a fence that impedes the diffusion of phosphatidylinositol 4,5-bisphosphate out of the forming phagosomes of macrophages. MBoC 22, 3498–3507 (2011).

34. M. E. Maxson, X. Naj, T. R. O’Meara, J. D. Plumb, L. E. Cowen, S. Grinstein, Integrin-based diffusion barrier separates membrane domains enabling the formation of microbiostatic frustrated phagosomes. eLife 7, e34798 (2018).

35. P. Beemiller, Y. Zhang, S. Mohan, E. Levinsohn, I. Gaeta, A. D. Hoppe, J. A. Swanson, A Cdc42 Activation Cycle Coordinated by PI 3-Kinase during Fc Receptor-mediated Phagocytosis. Mol Biol Cell 21, 470–480 (2010).

36. D. Cox, B. M. Dale, M. Kashiwada, C. D. Helgason, S. Greenberg, A Regulatory Role for Src Homology 2 Domain–Containing Inositol 5′-Phosphatase (Ship) in Phagocytosis Mediated by Fcγ Receptors and Complement Receptor 3 (α M β 2 ; Cd11b/Cd18). The Journal of Experimental Medicine 193, 61–72 (2001).

37. M. Bohdanowicz, S. Grinstein, Role of Phospholipids in Endocytosis, Phagocytosis, and Macropinocytosis. Physiological Reviews 93, 69–106 (2013).

38. G. D. Fairn, K. Ogata, R. J. Botelho, P. D. Stahl, R. A. Anderson, P. De Camilli, T. Meyer, S. Wodak, S. Grinstein, An electrostatic switch displaces phosphatidylinositol phosphate kinases from the membrane during phagocytosis. Journal of Cell Biology 187, 701–714 (2009).

39. R. J. Botelho, M. Teruel, R. Dierckman, R. Anderson, A. Wells, J. D. York, T. Meyer, S. Grinstein, Localized Biphasic Changes in Phosphatidylinositol-4,5-Bisphosphate at Sites of Phagocytosis. The Journal of Cell Biology 151, 1353–1368 (2000).

40. J. E. Ferrell, Perfect and Near-Perfect Adaptation in Cell Signaling. cels 2, 62–67 (2016).

41. P. Várnai, K. I. Rother, T. Balla, Phosphatidylinositol 3-Kinase-dependent Membrane Association of the Bruton’s Tyrosine Kinase Pleckstrin Homology Domain Visualized in Single Living Cells. Journal of Biological Chemistry 274, 10983–10989 (1999).

42. D. E. Schiff, J. Rae, T. R. Martin, B. H. Davis, J. T. Curnutte, Increased Phagocyte FcγRI Expression and Improved Fcγ-Receptor–Mediated Phagocytosis After In Vivo Recombinant Human Interferon-γ Treatment of Normal Human Subjects. Blood 90, 3187– 3194 (1997).

43. W. H. Fridman, I. Gresser, M. T. Bandu, M. Aguet, C. Neauport-Sautes, Interferon enhances the expression of Fc gamma receptors. The Journal of Immunology 124, 2436– 2441 (1980).

44. ImmGen Consortium, Open-source ImmGen: mononuclear phagocytes. Nat Immunol 17, 741 (2016).

45. D. J. Loegering, M. R. Lennartz, Signaling Pathways for Fcγ Receptor-Stimulated Tumor Necrosis Factor-α Secretion and Respiratory Burst in RAW 264.7 Macrophages. Inflammation 28, 23–31 (2004).

46. A. Ram, D. Murphy, N. DeCuzzi, M. Patankar, J. Hu, M. Pargett, J. G. Albeck, A guide to ERK dynamics, part 2: downstream decoding. Biochemical Journal 480, 1909–1928 (2023).

47. P. A. Gagliardi, O. Pertz, The mitogen-activated protein kinase network, wired to dynamically function at multiple scales. Current Opinion in Cell Biology 88, 102368 (2024).

48. S. D. M. Santos, P. J. Verveer, P. I. H. Bastiaens, Growth factor-induced MAPK network topology shapes Erk response determining PC-12 cell fate. Nat Cell Biol 9, 324–330 (2007).

49. C. J. Marshall, Specificity of receptor tyrosine kinase signaling: Transient versus sustained extracellular signal-regulated kinase activation. Cell 80, 179–185 (1995).

50. A. G. Goglia, M. Z. Wilson, S. G. Jena, J. Silbert, L. P. Basta, D. Devenport, J. E. Toettcher, A Live-Cell Screen for Altered Erk Dynamics Reveals Principles of Proliferative Control. Cell Systems 10, 240–253.e6 (2020).

51. S. Regot, J. J. Hughey, B. T. Bajar, S. Carrasco, M. W. Covert, High-sensitivity measurements of multiple kinase activities in live single cells. Cell 157, 1724–1734 (2014).

