## Supplemental Information for "Inhibitory Fc Receptor sets a time limit on macrophage response to IgG"

**The PDF file includes:**

Figs. S1 to S7
Tables S1 to S6

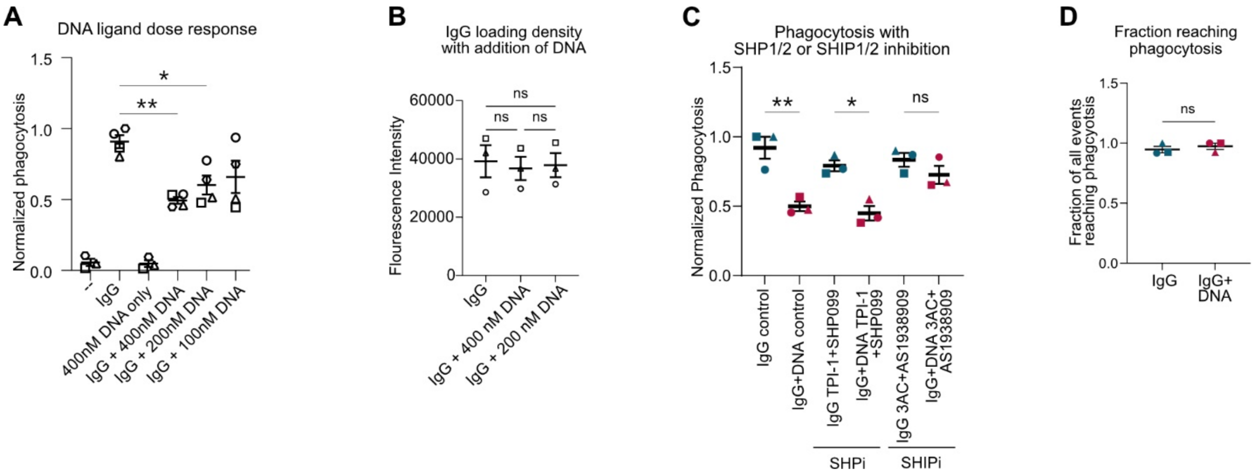


**Fig. S1. Related to figure 1. FcγRIIB DNA CAR inhibits phagocytosis via SHIP phosphatases.**

**(A)** FcγRIIB DNA CAR RAW264.7 macrophages were incubated with beads bound to neutravidin (--), IgG and neutravidin (IgG), or IgG, neutravidin and various concentrations of biotinylated ligand DNA (IgG+DNA). The average number of 5 µm diameter beads phagocytosed per macrophage was quantified by spinning disk confocal microscopy. **(B)** Graph shows loading density of atto488 labeled anti-biotin IgG on targets with the addition of ligand DNA. Atto488 fluorescence was measured by flow cytometry. **(C)** FcγRIIB DNA CAR macrophages were treated with SHP inhibitors in combination (TPI-1, SHP1 inhibitor and SHP099, SHP2 inhibitor) or SHIP inhibitors (3AC, SHIP1 inhibitor and AS1938909, SHIP2 inhibitor). After 12 hours of treatment, IgG or IgG+DNA beads were added and phagocytosis was measured by confocal microscopy. **(D)** Timelapse microscopy was used to quantify the fraction of phagocytic cups that extended beyond the macrophage cell cortex, indicating phagocytosis. Data points from replicates performed on the same day are denoted by symbol shape. Bars represent the mean +/- SEM. *indicates $p<0.05$, **indicates $p<0.01$ by ordinary one-way ANOVA with Dunnett's multiple comparisons test (A); ordinary one-way ANOVA with Tukey's multiple comparisons test (B); ordinary one-way ANOVA with Šídák's multiple comparisons test (C); or unpaired two-tailed t test (D).

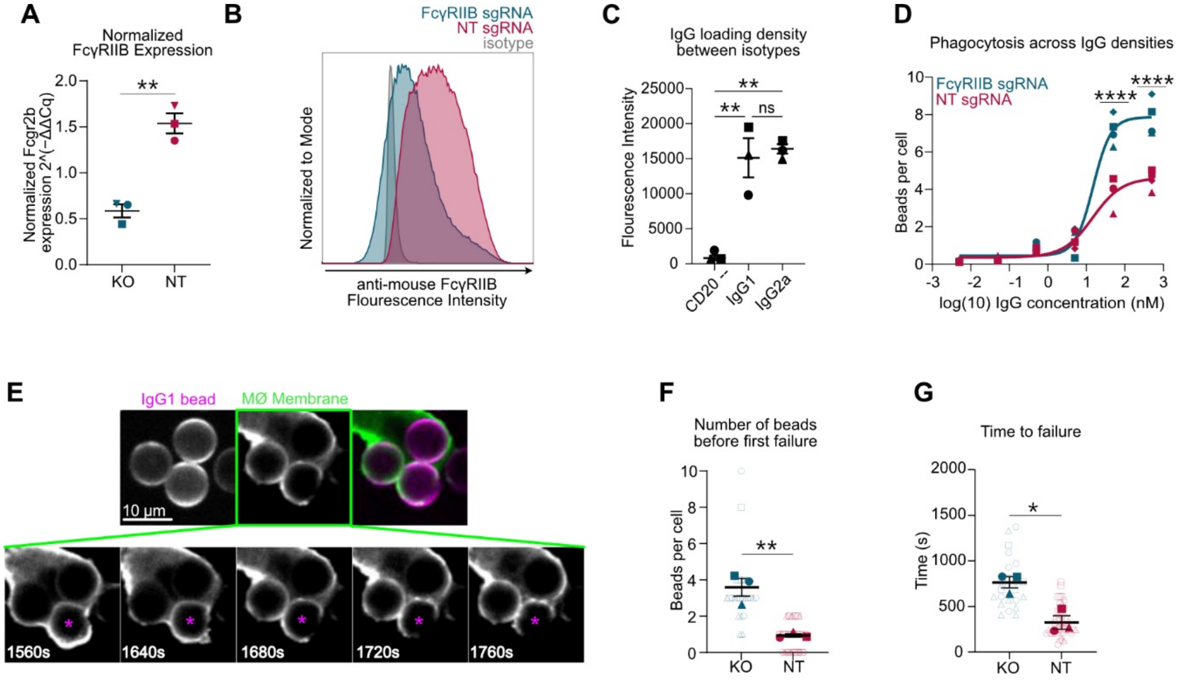


**Fig. S2. Related to Figure 1. FcγRIIB CRISPR knockout validation and impact on phagocytosis**

**(A)** *Fcgr2b* expression in HoxB8 macrophages infected with an sgRNA targeting FcγRIIB or a non-targeting guide was measured by qPCR. **(B)** Representative histograms show levels of FcγRIIB on the surface of HoxB8 macrophages infected with an sgRNA targeting FcγRIIB or a non-targeting guide measured by flow cytometry. **(C)** Graph shows loading density of mouse anti-CD20 IgG1 or IgG2a. Atto647 labeled anti-mouse antibody was used to detect anti-CD20 antibodies. Atto647 fluorescence was measured by flow cytometry. **(D)** Graph shows phagocytosis of 5 µm diameter beads coupled with increasing concentrations of IgG1 by HoxB8 macrophages infected with an sgRNA targeting FcγRIIB (blue) or a non-targeting guide (red). **(E)** spinning disk confocal images depict a HoxB8 NT sgRNA infected macrophage expressing a membrane marker (GFP-CAAX; green) failing to phagocytose a bead covered in a supported lipid bilayer (atto647; magenta) conjugated to IgG1. **(F-G)** Timelapse microscopy was used to quantify the number of beads successfully engulfed before the first observed failure (F) and time between initiating and failing phagocytosis (G). Solid data points represent the mean of an independent experiment, while unfilled data points represent individual cell measurements. Data points from replicates performed on the same day are denoted by symbol shape. In all graphs, bars represent the mean +/- SEM. *indicates $p<0.05$, **indicates $p<0.01$, ***indicates $p<0.001$ by ordinary one-way ANOVA with Tukey's multiple comparisons test (C); two-way ANOVA with Tukey's multiple comparisons test (D) unpaired two-tailed t test (A, F, G).

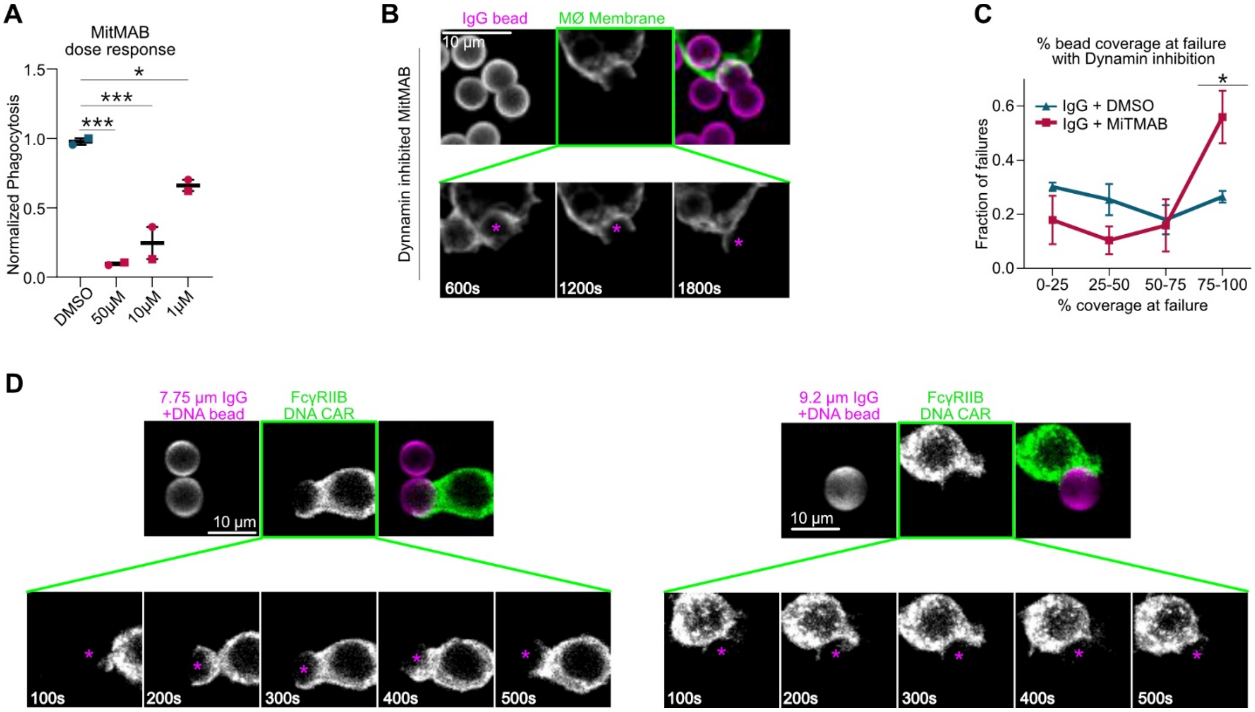


**Fig. S3. Related to figure 1. Dynamin regulates phagocytic cup closure**

**(A)** RAW264.7 macrophages were treated with MiTMAB to inhibit dynamin. Phagocytosis of IgG opsonized beads was measured by confocal microscopy. **(B)** spinning disk confocal images of a RAW264.7 macrophage stained with a membrane marker (CellTrace green) treated with 1uM MiTMAB failing to phagocytose a supported lipid bilayer coated bead (atto647; magenta) opsonized with anti-biotin IgG. **(C)** Graph shows the maximum percentage of the target covered by the macrophage membrane for IgG (red) or IgG+DNA (blue) targets after treatment with Dynamin inhibitor MiTMAB. **(D)** spinning disk confocal images of a macrophage expressing the DNA FcγRIIB CAR (mScarlet; green) failing to phagocytose a 7.75 µm or 9.2 µm glass bead coated in a supported lipid bilayer (atto647; magenta) opsonized with anti-biotin IgG and ligand DNA. In A, points represent independent replicates. In C, points represent the mean of 3 independent experiments. Bars represent the mean +/- SEM. *indicates $p<0.05$, ***indicates $p<0.001$ by ordinary one-way ANOVA with Šídák's multiple comparisons test (A); and two-way ANOVA with Šídák's multiple comparisons test (C)

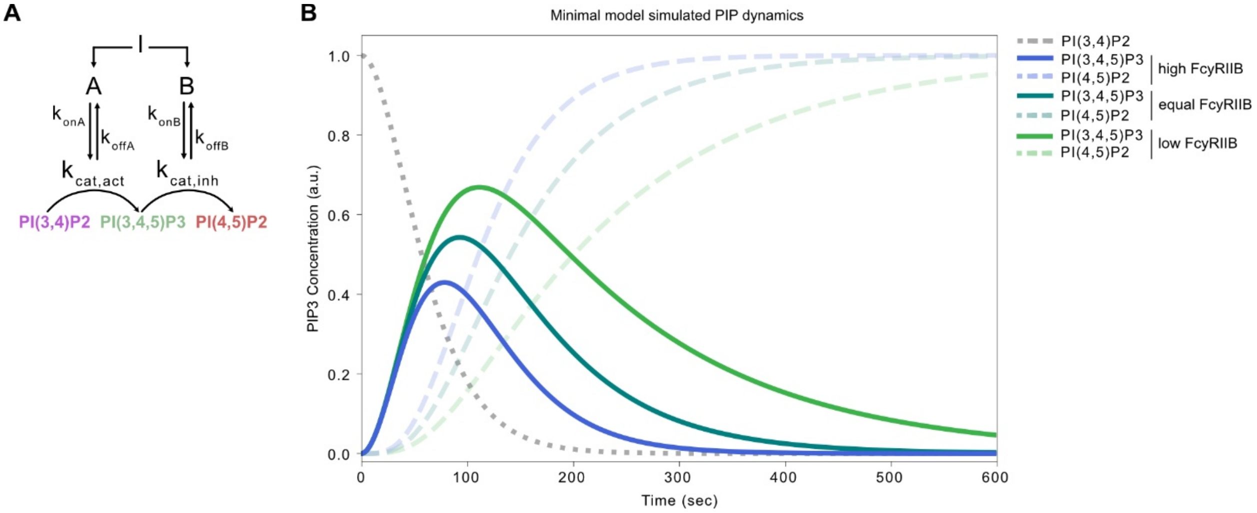


**Fig. S4. Related to figure 2. Minimalistic model of PIP3 dynamics during IgG binding.**
**(A)** Summary schematic shows the parameters incorporated into our minimal model. Specific values are in Table S5. **(B)** Graphs display PIP dynamics with varying levels of inhibitory input (B). As indicated in Table S5, amount of B is titrated through multiplier (ρB).

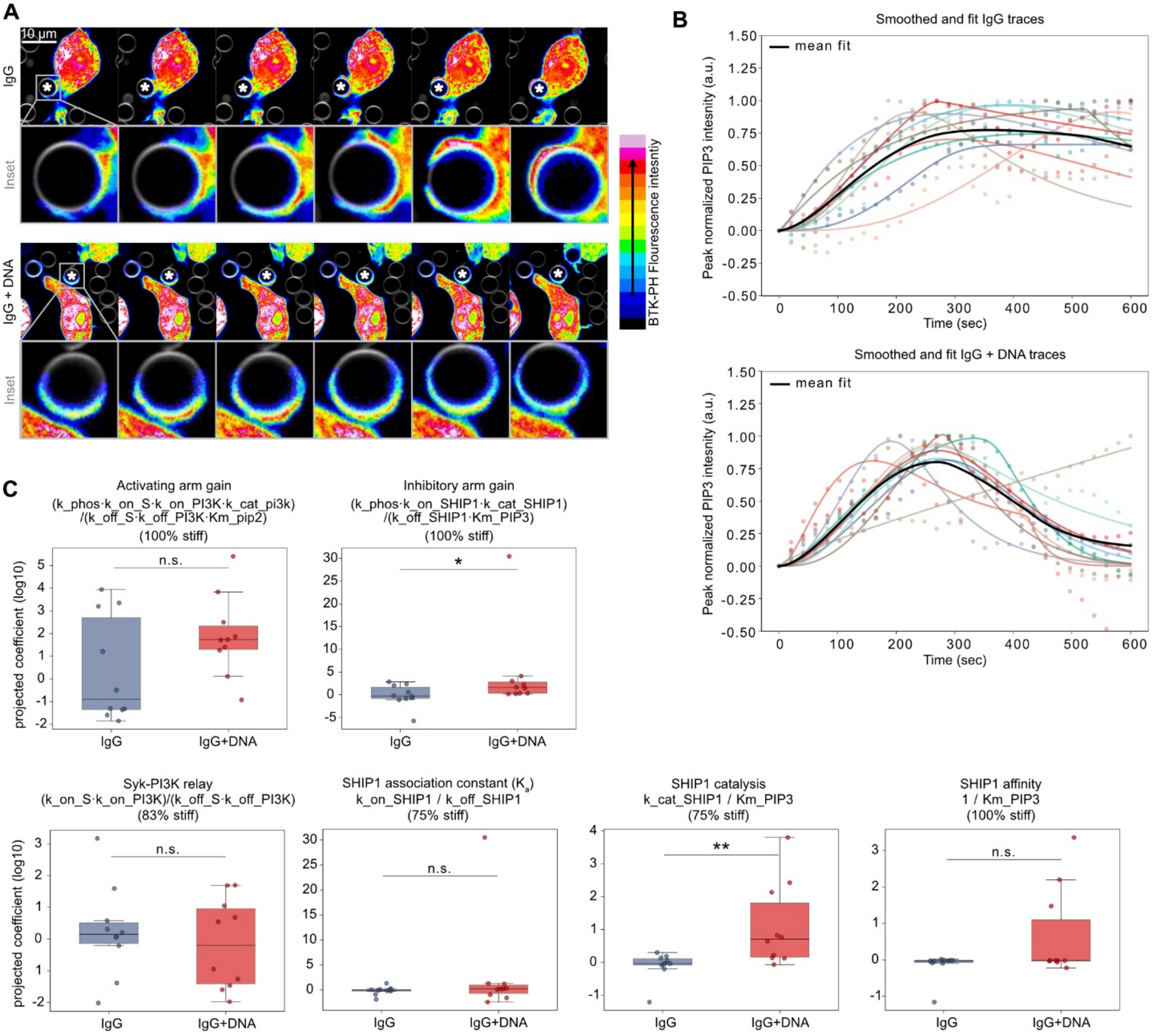


**Fig. S5. Related to figure 2. In silico model fit to measured PIP3 traces.**

**(A)** spinning disk confocal images show PIP3 reporter (BTK-PH-GFP; 16 color map) localization in FcγRIIB DNA CAR macrophages during phagocytosis of beads (atto647; grey) conjugated with IgG only (top) or IgG and DNA (bottom). **(B)** Cell-by-cell model fits to single-cell PIP3 traces at the phagocytic cup under IgG stimulation (left) and IgG + DNA (right). Fits were performed on baseline-subtracted traces. Points show single-cell measurements and solid lines show the corresponding least-squares ordinary differential equation solutions, color-matched by cell. Black lines indicate the mean of the ten single-cell fit trajectories. **(C)** Condition-level differences in identifiable parameter combinations (≥75% within the stiff subspace) between IgG and IgG + DNA. Per-cell values were computed from each cell's fitted rate constants (log10 units); boxes show median and interquartile range, whiskers 1.5× IQR. Inhibitory-arm combinations were compared by one-sided and activating-arm controls by two-sided Mann-Whitney U test. * $p < 0.05$, ** $p < 0.01$.

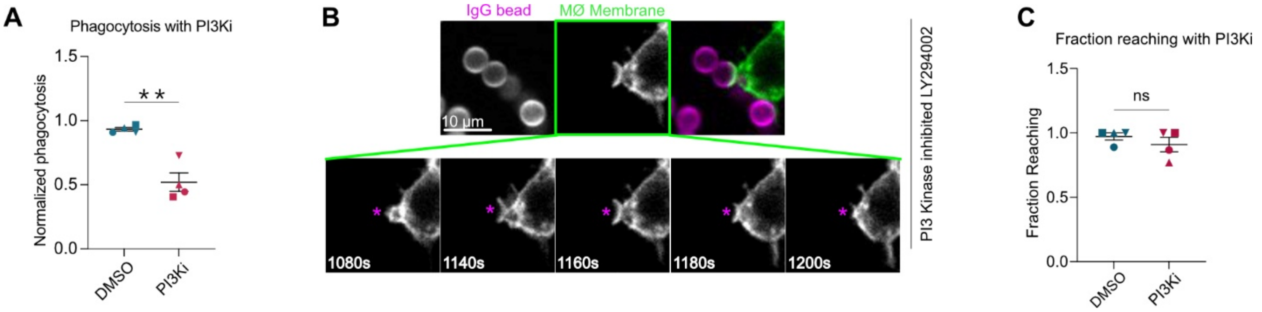


**Fig. S6. related to figure 3. PI3K inhibition reduces initiation of phagocytosis.**

**(A)** RAW264.7 macrophages were incubated with IgG beads with (red) or without (blue) pretreatment with PI3K inhibitor. Each data point represents the mean of 2 technical replicates, normalized to the highest phagocytosis level that day (n=50-150 cells per technical replicate) and are independent experiments. **(B)** spinning disk confocal images of a RAW264.7 macrophage expressing CAAX membrane marker (mCherry; green) treated with PI3K inhibitor failing to phagocytose a 5 µm glass bead coated in a supported lipid bilayer (atto647; magenta) opsonized with anti-biotin IgG. **(C)** Timelapse imaging was used to quantify the fraction of phagocytosis events that were qualified as reaching, as opposed to sinking phagocytosis. Each data point represents the mean of 10-50 bead contacts and is an independent experiment. Data points from replicates performed on the same day are denoted by symbol shape. **indicates $p<0.01$ by unpaired two tailed t test.

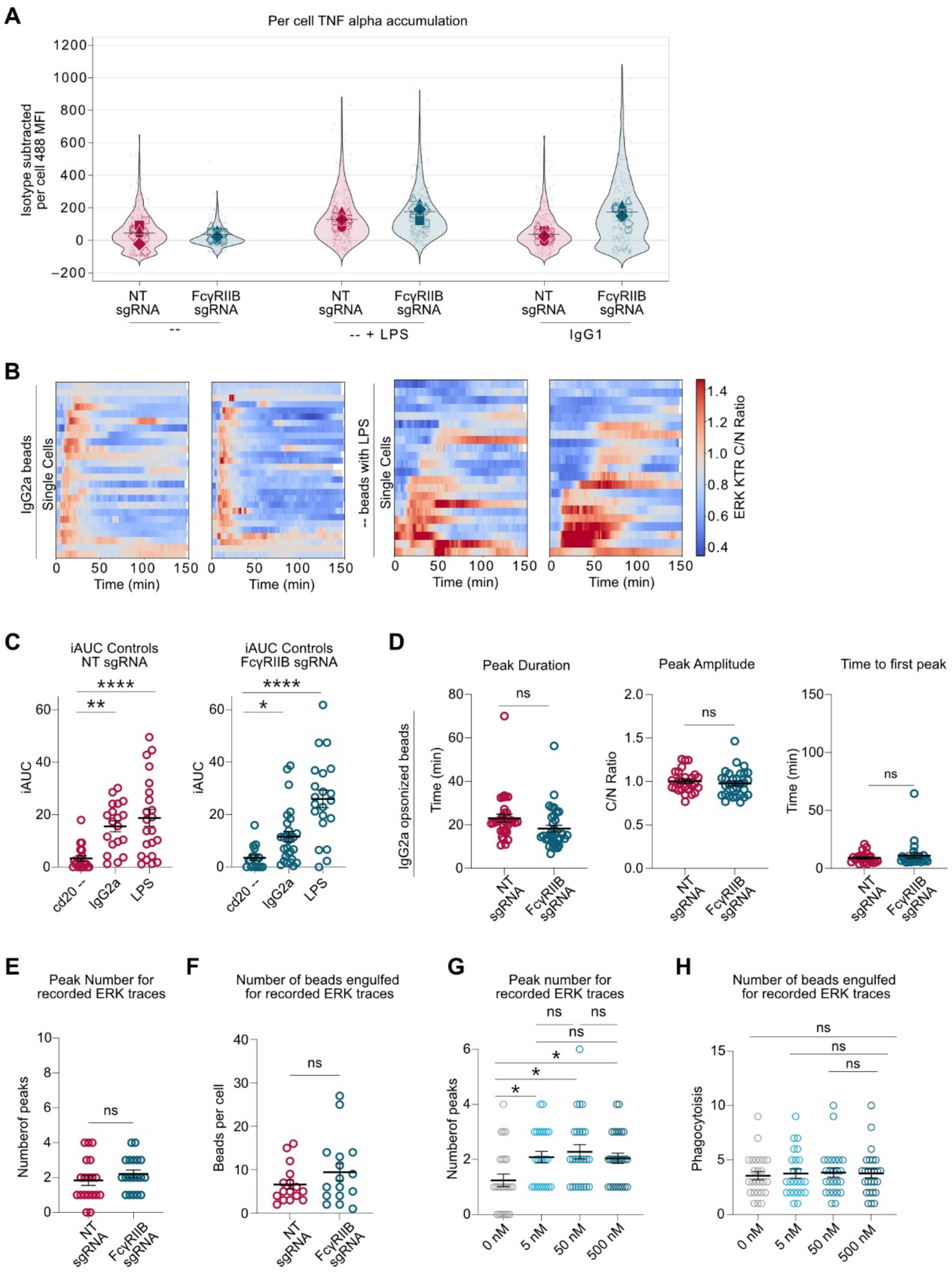


**Fig. S7. related to figure 4. TNFα and ERK response to IgG**

**(A)** HoxB8 macrophages infected with FcγRIIB KO or NT sgRNA were incubated with bead targets (unopsonized, unopsonized with LPS, or opsonized with IgG1) for 30 minutes before a 4 hour incubation with Brefeldin A to trap cytokines intracellularly for later staining and confocal imaging. Violin plots show per cell TNFα staining (antiTNFα-488). **(B)** Heatmaps depict C/N ratio over time after phagocytosis of IgG2a opsonized beads (left) or unopsonized beads with 10 ng/ml LPS (right). **(C)** iAUC of ERK KTR traces after engulfment of unopsonized CD20-bound beads with and without LPS, or IgG2a opsonized beads in HoxB8 macrophages infected with FcγRIIB KO sgRNA or NT sgRNA. LPS was added to imaging wells immediately preceding addition of beads and the start of imaging. **(D)** Graphs display peak duration, peak amplitude, and time to first peak for C/N ratio traces in HoxB8 macrophages infected with FcγRIIB KO or NT sgRNA after engulfment of IgG2a beads. **(E)** Graph displays the number of peaks for traces displayed in Fig. 4D. **(F)** Graph displays the number of beads engulfed for corresponding traces displayed in Fig. 4D. Our inclusion criteria was designed to target cells that phagocytosed a similar number of beads. **(G)** Graph displays the number of peaks for traces displayed in Fig. 4F. **(H)** Graph displays the number of beads engulfed for corresponding traces displayed in Fig. 4F. For A, solid points represent the mean of independent experiments, unfilled data points represent technical replicates performed on the same day, and small points represent individual cells. For other graphs, each data point represents a different cell, collected across 3 independent experiments. In all graphs, bars represent the mean +/- SEM. *indicates $p<0.05$, **indicates $p<0.01$, ***indicates $p<0.001$ by two-way ANOVA with Tukey's multiple comparisons test (C), Welch's unpaired t test (D-F), or Kruskal-Wallis test with Dunn's multiple comparisons test (G,H).

**Table S1. Expressed reporters, constructs, and oligos used for qPCR and DNA receptor-ligand pairs.**

| Constructs | |
|---|---|
| **Name and source** | **Description** |
| pHR FcγR DNA CAR - Addgene #176611 | In pHR vector. Signal peptide: (MQSGTHWRVLGLCLLSVGVWGQD) derived from CD3ε Extracellular: HA tag,a linker (LPETGGGGGG), SNAPf (from the pSNAPf plasmid, New England Biolabs) Linker: GGSGGSGGS, TM and intracellular: (aa 19–86) of the Fcγ-chain UniProtKB – P20491 (FCERG_MOUSE) with aa25 mutated from C to A linker: GSGS, Fluorophore: mGFP. |
| pHR FcγRIIB DNA CAR - this paper | In pHR vector. Signal peptide: (MQSGTHWRVLGLCLLSVGVWGQD) derived from CD3ε. Extracellular: HA tag plus a linker (LPETGGGGGG), HaloTag (from the HaloTag plasmid, New England Biolabs). Linker: GGSGGSGGS. TM and intracellular: TM and intracellular domain (aa211-283) of mouse Fcgr2b myeloid isoform B2 UniProtKB – P08101-2 (FCGR2_MOUSE), Linker: TSPVAT. Fluorophore: mScarlet. |
| pHR BTK-PH-GFP - this paper | BTK-PH-GFP PIP3 biosensor from (Várnai et al., 1999; Marshall et al., 2001) was cloned into the pHR vector. |
| Fcgr2b and nontargeting sgRNA plasmids - this paper | Third-generation lentiviral sgRNA plasmids (8,577 bp) built on the lenti-sgRNA blast backbone (Stringer et al., 2019), expressing an S. pyogenes sgRNA from a U6 promoter and blasticidin resistance from an EF-1α promoter. The Fcgr2b targeting construct carries 5′-GCCGTTCCTACTGATCCCCA-3′, for mouse Fcgr2b (UniProtKB P08101, FCGR2_MOUSE). The non-targeting control construct carries 5′-GCGAGGTATTCGGCTCCGCG-3′, which has no match in the mouse or human genome. Plasmids were cloned in accordance with an established protocol by Zhang et al. described in the Addgene repository for the lentiviral backbone (Addgene #104993). |
| pHR ERK KTR mClover - addgene #59150 | — |
| pHR GFP CAAX - addgene #113020 (Morrissey et al., 2018) | — |

| | |
|---|---|
| pHR H2B mCh - addgene #89766 | — |
| **Oligos** | |
| **qPCR** | |
| **Gapdh** | Primer 1: 5'-AATGGTGAAGGTCGGTGTG-3' |
| | Primer 2: 5'-GTGGAGTCATACTGGAACATGTAG-3' |
| **Fcgr2b** | Primer 1: 5'-CAGTGGAGAATATCGGTGTCAA-3' |
| | Primer 2: 5'-CCCCTTCCAGAAACACCAG-3' |
| DNA based receptor-ligand pairs | |
| **Activating FcR DNA CAR** | Receptor strand: 5’ - BG-/5AmMC6/AATATGATGTATGTGG - 3' |
| | Ligand strand: 5' - /5BiosG/TTTT-TTTCATACATCATATT-3' |
| **FcγRIIB DNA CAR** | Receptor strand: 5’ - HALO-/5AmMC6/AAACTTCGACATGATA - 3’ |
| | Ligand strand: 5’ - /5BiosG/TATCATGTCGAAGTTT - 3’ |

**Table S2. Coverage and surface area of the phagocytic cup at failure, by bead diameter.**

| Bead diameter (µm) | Coverage at failure (%) | Surface area covered at failure (µm²) |
|---|---|---|
| 1.86 | 540–723 | 58–79 |
| 4.3 | 100–135 | 58–79 |
| 5.0 | 75–100 | 58–79 |
| 7.75 | 31–41 | 58–79 |
| 9.2 | 22–29 | 58–79 |

Total surface area of a bead: $SA_{total} = \pi d^2$

Reference area (from 5 µm bead, 75–100% bin): $SA_{covered} = (\%coverage / 100) \times \pi(5)^2 =$ 58–79 $\mu m^2$

Applied to other diameters: $\%coverage = (58\text{–}79\ \mu m^2) / (\pi d^2) \times 100$

**Table S3. ODEs for full model.**

| Equation | Description |
|---|---|
| **Receptor–antibody binding and FcγR motif phosphorylation** | |
| $d(FcR\_A)/dt = -k_{bind,A} \cdot Ab\_A \cdot FcR\_A + k_{unbind,A} \cdot FcR\_Ab\_A$ | Free activating FcγR |
| $d(FcR\_Ab\_A)/dt = k_{bind,A} \cdot Ab\_A \cdot FcR\_A - k_{unbind,A} \cdot FcR\_Ab\_A - k_{phos} \cdot FcR\_Ab\_A$ | Ab-bound activating FcγR |
| $d(FcRp\_A)/dt = k_{phos} \cdot FcR\_Ab\_A$ | Phosphorylated activating FcγR (pITAM) |
| $d(Ab\_A)/dt = -k_{bind,A} \cdot Ab\_A \cdot FcR\_A + k_{unbind,A} \cdot FcR\_Ab\_A$ | Free antibody for activating arm |
| $d(FcR\_B)/dt = -k_{bind,B} \cdot Ab\_B \cdot FcR\_B + k_{unbind,B} \cdot FcR\_Ab\_B$ | Free FcγRIIB |
| $d(FcR\_Ab\_B)/dt = k_{bind,B} \cdot Ab\_B \cdot FcR\_B - k_{unbind,B} \cdot FcR\_Ab\_B - k_{phos} \cdot FcR\_Ab\_B$ | Ab-bound FcγRIIB |
| $d(FcRp\_B)/dt = k_{phos} \cdot FcR\_Ab\_B$ | Phosphorylated FcγRIIB (pITIM) |
| $d(Ab\_B)/dt = -k_{bind,B} \cdot Ab\_B \cdot FcR\_B + k_{unbind,B} \cdot FcR\_Ab\_B$ | Free antibody for inhibitory arm |
| **Syk and PI3K recruitment** | |
| $dS/dt = k_{on,S} \cdot FcRp\_A - k_{off,S} \cdot S$ | Active Syk; pseudo-first-order recruitment to pITAM |
| $d(PI3K)/dt = k_{on,PI3K} \cdot S - k_{off,PI3K} \cdot PI3K$ | Membrane-recruited PI3K |
| **SHIP1 recruitment** | |
| $d(SHIP1)/dt = k_{on,SHIP1} \cdot FcRp\_B - k_{off,SHIP1} \cdot SHIP1$ | pITIM-recruited SHIP1; mirrors PI3K recruitment by Syk |
| **PIP lipid dynamics** | |
| $d(PIP2)/dt = -R_{gen}$ | $PI(4,5)P_2$ consumption by PI3K |
| $d(PIP3)/dt = R_{gen} - R_{dep}$ | $PI(3,4,5)P_3$ net flux |

| | |
|---|---|
| $d(PI34P2)/dt = R_{dep}$ | $PI(3,4)P_2$ production (SHIP1 product) |
| **Rate definitions** | |
| $R_{gen} = k_{cat,PI3K} \cdot PI3K \cdot (PIP2 / (K_m PIP2 + PIP2))$ | PI3K-catalyzed $PI(3,4,5)P_3$ generation (Michaelis–Menten) |
| $R_{dep} = k_{cat,SHIP1} \cdot SHIP1 \cdot (PIP3 / (K_m PIP3 + PIP3))$ | SHIP1-mediated $PI(3,4,5)P_3$ dephosphorylation (Michaelis–Menten) |

**Table S4. Full model parameters and values.**

Bimolecular on-rates are listed as reported values ($M^{-1}s^{-1}$); multiplied by Cref = 1 μM in the ODE system.

| Parameter | Description | Value | Units | Source | Notes |
|---|---|---|---|---|---|
| **$K_1$** $k_{bind,A}$ | $Ab_A$ association with FcγRIIIA | $6.51 \times 10^3$ | $M^{-1}s^{-1}$ | Li et al., 2007 (Li et al., 2007) | — |
| **$K_2$** $k_{unbind,A}$ | $Ab_A$ dissociation from FcγRIIIA | $4.71 \times 10^{-3}$ | $s^{-1}$ | Li et al., 2007 (Li et al., 2007) | Derived from $K_D$ = 0.724 μM |
| **$K_3$** $k_{bind,B}$ | $Ab_B$ association with FcγRIIB | $3.80 \times 10^5$ | $M^{-1}s^{-1}$ | Maenaka et al., 2001 (Maenaka et al., 2001) | — |
| **$K_4$** $k_{unbind,B}$ | $Ab_B$ dissociation from FcγRIIB | 0.88 | $s^{-1}$ | Maenaka et al., 2001 (Maenaka et al., 2001) | Derived from $K_D$ = 2.32 μM |
| **$K_5$** $k_{phos}$ | SFK phosphorylation of ITAM/ITIM | 0.15 | $s^{-1}$ | Barua et al., 2012; Suter et al, 2021 (Barua et al., 2012; Suter et al., 2021) | — |
| **$K_6$** $k_{on,S}$ | Syk recruitment to pITAM | $1.5 \times 10^5$ | $M^{-1}s^{-1}$ | Faeder et al., 2003; Barua et al., 2012 (Faeder et al., 2003; Barua et al., 2012) | — |
| **$K_7$** $k_{off,S}$ | Syk dissociation from pITAM | 0.13 | $s^{-1}$ | Faeder et al., 2003 (Faeder et al., 2003) | — |
| **$K_8$** $k_{on,PI3K}$ | PI3K recruitment to active Syk | $3.34 \times 10^6$ | $M^{-1}s^{-1}$ | Panayotou et al., 1993 (Panayotou et al., 1993) | — |
| **$K_9$** $k_{off,PI3K}$ | PI3K dissociation from Syk | 1.002 | $s^{-1}$ | Ladbury et al., 1995 (Ladbury et al., 1995) | Derived from $K_D$ = 300 nM |
| **$K_{12}$** $k_{cat,PI3K}$ | $PI(4,5)P_2 \rightarrow PI(3,4,5)P_3$ turnover | 0.975 | $s^{-1}$ | Huang et al., 2011 (Huang et al., 2011) | Derived from $V_{max}$ = 0.28 pmol $ng^{-1}min^{-1}$ |

| | | | | | |
|---|---|---|---|---|---|
| $K_m$PIP2 | Michaelis constant for PI3K | 1.80 | μM | Maheshwari et al., 2017 (Maheshwari et al., 2017) | — |
| **$K_{10}$** $k_{on,SHIP1}$ | SHIP1 recruitment to pITIM | 1.05 × $10^6$ | $M^{-1}s^{-1}$ | Zhang et al., 2009 (Zhang et al., 2009) | — |
| **$K_{11}$** $k_{off,SHIP1}$ | SHIP1 dissociation from pITIM | 0.29 | $s^{-1}$ | Zhang et al., 2009 (Zhang et al., 2009) | Derived from $K_D$ = 0.40 μM |
| **$K_{13}$** $k_{cat,SHIP1}$ | $PI(3,4,5)P_3 \rightarrow PI(3,4)P_2$ turnover | 8.0 | $s^{-1}$ | Waddell et al., 2023 (Waddell et al., 2023) | — |
| $K_m$PIP3 | Michaelis constant for SHIP1 | 94.0 | μM | Waddell et al., 2023 (Waddell et al., 2023) | — |

**Table S5. Minimal model parameters and equations.**

All intrinsic rate constants are identical between the activating and inhibitory arms. Only the FcγRIIB expression multiplier (ρB) differs across conditions; all parameter values are dimensionless (arbitrary units). The activator (A) and inhibitor (B) arms follow first-order recruitment and decay, and lipid conversion is mass-action. ρB scales only the inhibitor recruitment term, representing variation in FcγRIIB surface density.

| Parameter | Description | Value | Units |
|---|---|---|---|
| **Model Parameters** | | | |
| $k_{on,A}$ | Activator recruitment rate | 0.020 | a.u. |
| $k_{off,A}$ | Activator decay rate | 0.020 | a.u. |
| $k_{on,B}$ | Inhibitor recruitment rate | 0.020 | a.u. |
| $k_{off,B}$ | Inhibitor decay rate | 0.020 | a.u. |
| $k_{cat,act}$ | $PI(4,5)P_2 \rightarrow PI(3,4,5)P_3$ catalytic rate | 0.030 | a.u. |
| $k_{cat,inh}$ | $PI(3,4,5)P_3 \rightarrow PI(3,4)P_2$ catalytic rate | 0.030 | a.u. |
| $\rho_B$ | FcγRIIB expression multiplier | 1.8 / 1.0 / 0.5 | dimensionless |
| I(t) | IgG input (step function) | 1.0 for $t > 0$ | a.u. |
| **Rate Equations** | | | |
| Activator (A) | $dA/dt = k_{on,A} \cdot I(t) - k_{off,A} \cdot A$ | | |
| Inhibitor (B) | $dB/dt = \rho_B \cdot k_{on,B} \cdot I(t) - k_{off,B} \cdot B$ | | |
| $PI(4,5)P_2$ | $d[PIP_2]/dt = -k_{cat,act} \cdot A \cdot [PIP_2]$ | | |
| $PI(3,4,5)P_3$ | $d[PIP_3]/dt = k_{cat,act} \cdot A \cdot [PIP_2] - k_{cat,inh} \cdot B \cdot [PIP_3]$ | | |
| $PI(3,4)P_2$ | $d[PIP_{2,34}]/dt = k_{cat,inh} \cdot B \cdot [PIP_3]$ | | |

**Table S6. FcγR expression and activating-to-inhibitory ratio across tissue macrophage populations.** Values are RNA-seq read counts. Ratio = sum of activating receptors (FcγRI, III, IV) ÷ FcγRIIB.

| | **Spleen macrophages (GSM3455384)** | **Kupffer cells (GSM3455189)** | **Embryonic macrophages (GSM3455048)** | **Peritoneal macrophages (GSM3455034)** | **Thymic macrophages (GSM3455075)** |
|---|---|---|---|---|---|
| Fcgr1 | 402 | 1,458 | 605 | 207 | 720 |
| Fcgr2b | 20 | 216 | 135 | 1,460 | 3,483 |
| Fcgr3 | 1,321 | 2,713 | 1,690 | 5,852 | 1,540 |
| Fcgr4 | 1,112 | 3,133 | 38 | 254 | 92 |
| **Activating: Inhibitory ratio** | **141.75** | **33.81** | **17.28** | **4.32** | **0.68** |
